# A BioBricks^®^ toolbox for multiplexed metabolic engineering of central carbon metabolism in the tetracenomycin pathway

**DOI:** 10.1101/2021.07.23.453519

**Authors:** Jennifer T. Nguyen, Kennedy K. Riebschleger, Katelyn V. Brown, Nina M. Gorgijevska, S. Eric Nybo

## Abstract

The tetracenomycins are aromatic anticancer polyketides that inhibit peptide translation via binding to the large ribosomal subunit. Here, we expressed the elloramycin biosynthetic gene cluster in the heterologous host *Streptomyces coelicolor* M1146 to facilitate the downstream production of tetracenomycin analogs. We developed a BioBricks® genetic toolbox of genetic parts for substrate precursor engineering in *S. coelicolor* M1146::cos16F4iE. We cloned a series of integrating vectors based on the VWB, TG1, and SV1 integrase systems to interrogate gene expression in the chromosome. We genetically engineered three separate genetic constructs to modulate tetracenomycin biosynthesis: 1) the *vhb* hemoglobin from obligate aerobe *Vitreoscilla stercoraria* to improve oxygen utilization; (2) the *accA2BE* acetyl-CoA carboxylase to enhance condensation of malonyl-CoA; (3) lastly, the *sco6196* acyltransferase, which is a “metabolic regulatory switch” responsible for mobilizing triacylglycerols to β-oxidation machinery for acetyl-CoA. In addition, we engineered the *tcmO* 8-*O-*methyltransferase and newly identified *tcmD* 12-*O*-methyltransferase from *Amycolatopsis sp*. A23 to generate tetracenomycins C and X. We also co-expressed the *tcmO* methyltransferase with oxygenase *urdE* to generate the analog 6-hydroxy-tetracenomycin C. Altogether, this system is compatible with the BioBricks® [RFC 10] cloning standard for the co-expression of multiple gene sets for metabolic engineering of *Streptomyces coelicolor* M1146::cos16F4iE.

## Introduction

The tetracenomycins are a family of aromatic polyketides produced by *Streptomyces glaucescens* GLA.0 and *Streptomyces olivaceus* Tü2353, respectively ^[1,2]^. 8-demethyltetracenomycin C (8-DMTC, **1**), tetracenomycin C (**2**), tetracenomycin X (**3**), 6-hydroxy-tetracenomycin C (4), and elloramycin (**5**) are structurally representative compounds from this family (Figure 1). **1** - **5** exhibit antibacterial activity against gram-positive microorganisms and anticancer activity against a variety of mammalian cancer cell lines, though **2** and **3** are the most potent compounds described to date ^[3,4]^. The tetracenomycins were previously thought to exhibit a mechanism of action like the anthracyclines, namely, binding to DNA topoisomerase II and induction of DNA damage ^[3]^. Recently, Osterman and coworkers demonstrated that **3** does not induce DNA damage, but rather it inhibits peptide translation via binding to the large ribosomal subunit polypeptide exit channel ^[5]^. Impressively, Osterman et al. demonstrated that **3** binds to the same binding site within the exit channel of the *E. coli* 50S ribosomal subunit and the *H. sapiens* 60S ribosomal subunit, a stunning display of evolutionarily conserved molecular recognition that accounts for the cytotoxic activities of the TCMs ^[5]^. This sets the stage for the development of tetracenomycin analogs with improved potency and anticancer activity.

**Figure 1.**
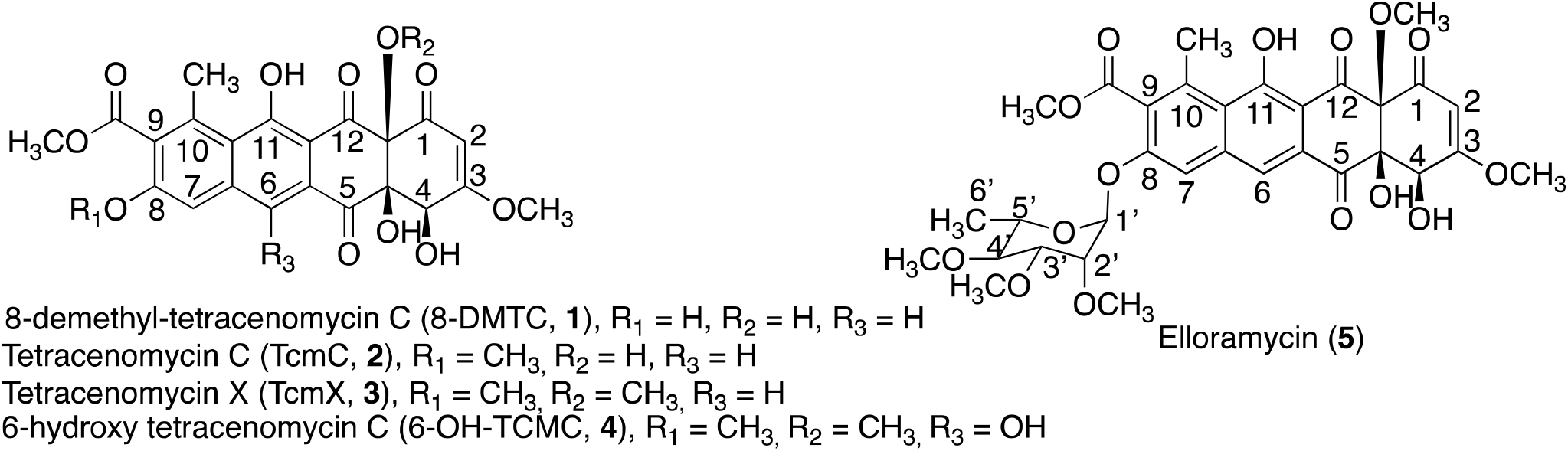
Structures of the antitumor tetracenomycins.

The gene cluster for *Streptomyces glaucescens* GLA.0 was previously sequenced, revealing the full biosynthetic gene cluster for **1**, spectinomycin, and acarbose ^[6]^. The biosynthesis of **1** has been studied extensively for the past three decades and has served as a model system for understanding type II polyketide biosynthesis (Figure 2) ^[7–11]^. Decker et al. previously isolated the gene cluster for elloramycin biosynthesis and cloned it onto cosmid cos16F4 ^[12]^. Heterologous expression of cosmid cos16F4 resulted in heterologous production of **1** and production of penultimate intermediate tetracenomycin A2 (Figure 2). Cosmid cos16F4 was also discovered to encode the *elmGT* gene, which encodes the glycosyltransferase responsible for the transfer of TDP-L-rhamnose to the 8-position of **1** ^[13]^. ElmGT has been shown to exhibit donor substrate flexibility towards >20 TDP-deoxysugar donors ^[14–20]^. Therefore, ElmGT and the cos16F4 heterologous expression system are significant tools for the generation of a library of tetracenomycin analogs. These tetracenomycin analogs will be instrumental in investigating the anticancer mechanism of action activity for this class of compounds and the role of the carbohydrate moiety in binding to the large mammalian ribosomal subunit.

**Figure 2.**
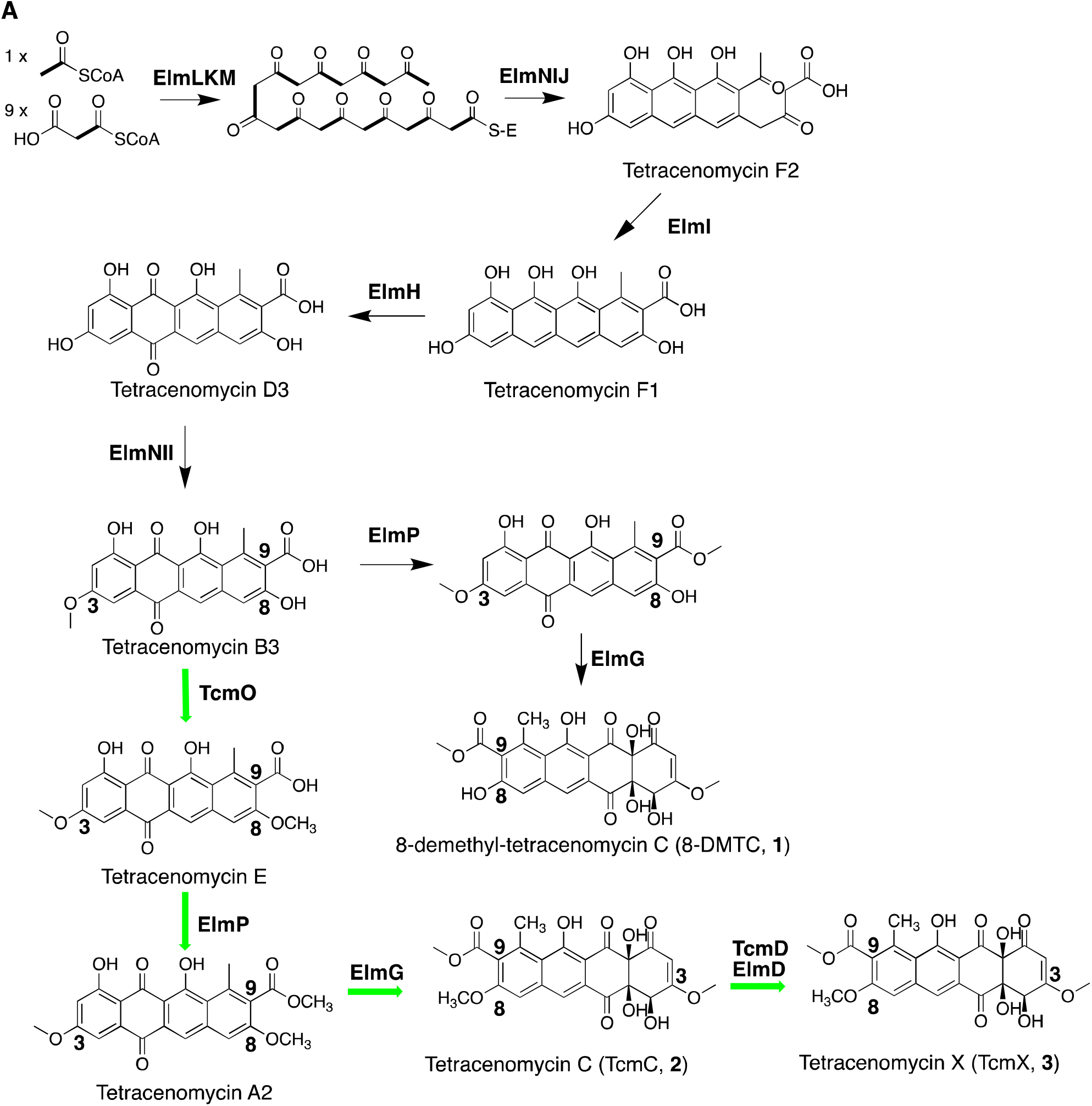
Proposed biosynthetic steps for the biosynthesis of the tetracenomycins. The native **1** biosynthetic pathway is indicated in black arrows, whereas the engineered tetracenomycin bypass pathway is indicated with bold green arrows.

The heterologous expression of extrachromosomal sequences in *Streptomyces spp*. Is subject to genetic instability ^[21]^. As an alternative approach, cloning of genes via the well-characterized actinophage integrases (e.g. fC31, fBT1, SV1, TG1, SAM2, VWB) into *attB* sites in the *Streptomyces* spp. chromosome can result in the stable incorporation of heterologously expressed genes ^[22]^. Therefore, we incorporated the elloramycin biosynthetic gene cluster encoded on the C31-integrating cassette cos16F4iE into the genome of the superhost *Streptomyces coelicolor* M1146. *Streptomyces coelicolor* M1146 has been genome minimalized for the removal of the actinorhodin, undecylprodigiosin, coelimycin P1, and the calcium-dependent antibiotic gene clusters ^[23]^. Therefore, *S. coelicolor* M1146 exhibits fungible metabolism that can be channeled towards the synthesis of natural products through the heterologous expression of biosynthetic gene clusters (BGCs).

In this work, we developed a BioBricks® [RFC 10] biosynthetic toolbox for the engineering of central carbon metabolism in elloramycin biosynthesis. First, we engineered the cos16F4iE cluster into *Streptomyces coelicolor* M1146 to generate an improved production host for tetracenomycins, as compared to the original *Streptomyces lividans* TK24 (cos16F4) host. In addition, we generated integrating plasmid cassettes based on plasmids pENSV1, pENTG1, and pOSV808 vectors to site-specifically introduce genes into the SV1, TG1, and VWB actinophage *attB* sites, respectively ^[24–27]^. Using these multiplexed integrating vectors, we engineered three different gene cassettes into different genomic loci to determine the optimal arrangement for enhancement of tetracenomycin production and biomass accumulation. First, we engineered the *vhb* hemoglobin gene from *Vitreoscilla stercoraria* to enhance aerobic respiration of *S. coelicolor* M1146::cos16F4iE in shake flasks ^[28,29]^. Secondly, we engineered the acetyl-CoA carboxylase *accA2BE* operon from *S. coelicolor* M145 under the control of the constitutive *ermE*p* promoter to enhance the production of malonyl-CoA ^[30]^. Thirdly, we engineered the *sco6196* acyltransferase, previously identified as a “metabolic switch” during the transition from triacylglycerol synthesis to polyketide biosynthesis in the stationary phase ^[31]^. Overexpression of the *accA2BE* operon and *sco6196* metabolic switches resulted in a 3-fold improvement in **1** production titer, as compared to the original production host. Finally, we utilized the improved production host expressing the *accA2BE* operon to engineer in the *tcmO* 8-*O*-methyltransferase and the *tcmD* 12-*O*-methyltransferase from *Amycolatopsis sp. A23* to biosynthesize tetracenomycin C and tetracenomycin X. To the best of our knowledge, this is the first report to describe the functional characterization of *tcmO* and *tcmD* in the biosynthesis of tetracenomycin X.

## Materials and Methods

### Bacterial strains and growth conditions

*E. coli* JM109 and *E. coli* ET12567 were grown at 37°C in LB broth or LB agar as previously described ^[32]^. *E. coli* JM109 was used for plasmid propagation and subcloning, while *E. coli* ET12567/pUZ8002 was used as the conjugation donor host for mobilizing expression vectors into *Streptomyces coelicolor* M1146 as previously described ^[33]^. (When appropriate, ampicillin (100 µg mL^-1^), kanamycin (25 µg mL^-1^), apramycin (25 µg mL^-1^), viomycin (30 µg mL^-1^), hygromycin (50 µg mL^-1^), and nalidixic acid (35 µg mL^-1^) were supplemented to media to select for recombinant microorganisms.

*Streptomyces coelicolor* M1146 and derivative strains were routinely maintained on Soya-Mannitol Flour (SFM) agar supplemented with 10 mM MgCl_2_ and International Streptomyces Project medium #4 (ISP4) (BD Difco) at 30°C as described previously ^[34]^. For liquid culturing, *Streptomyces coelicolor* M1146::cos16F4iE derivative strains were grown in TSB media for the production of seed culture and modified SG-TES liquid medium (soytone 10 g, glucose 20 g, yeast extract 5 g, TES free acid 5.73 g, CoCl_2_ 1 mg, per liter) ^[17]^. All media and reagents were purchased from Thermo-Fisher Scientific.

### General genetic manipulations

Routine genetic cloning and plasmid manipulation were carried out in *E. coli* JM109 (New England Biolabs). *E. coli* ET12567/pUZ8002 was used as the host for intergeneric conjugation with *Streptomyces coelicolor* as previously described ^[34]^. *E. coli* chemically competent cells were prepared using the Mix and Go! *E. coli* Transformation Kit® (Zymo Research). *E. coli* was transformed with plasmid DNA via chemically competent heat-shock transformation as described previously ^[32]^. Plasmid DNA was isolated via the Wizard® *Plus* SV Minipreps DNA Purification System by following the manufacturer’s protocols (Promega). All molecular biology reagents and enzymes used for plasmid construction were purchased from New England Biolabs.

BioBricks® parts were constructed to adhere to the BioBrick RFC[10] standard as previously described ^[35]^. For the construction of BioBrick® vectors, plasmids pSB1A3-J04450, pSB1K3-J04450, pSB1C3-J04450, pSB1T3-J04450 were used as previously described ^[36]^. In brief, these vectors encode the BBa_J04450 red fluorescent protein (RFP) coding device, which consists of the LacI promoter, the B0034 strong ribosome binding site, the monomeric red fluorescent protein from *Discosoma striata* (mRFP1), and the B0015 transcriptional terminator. *E. coli* JM109 strains transformed with these vectors develop a red color after approximately 18 hours, which indicates the presence of the RFP coding device. pSB1A3-J04450, pSB1K3-J04450, pSB1C3-J04450, pSB1T3-J04450 were restriction digested with *EcoRI*/*PstI* and treated with recombinant shrimp alkaline phosphatase, and used directly for subcloning without gel purification. In general, “5’-BioBricks® parts” were digested with *EcoRI/SpeI*, “3’-BioBricks® parts” were digested with *XbaI/PstI*, and these were spliced together in a three-way ligation (3A cloning) into a destination vector part with a different drug resistance marker ^[36]^. BioBricks® parts were spliced into the digested vectors, transformed into *E. coli* JM109 competent cells, and plated on LB agar with antibiotics. In a manner analogous to blue-white colony screening, colonies that turned white contained the genes of interest, whereas colonies that turned red were still expressing the RFP-coding device. Gene cassettes were assembled in pSB1A3, pSB1C3, pSB1K3, or pSB1T3 before restriction digestion with *EcoRI*/*PstI* and ligation to the same sites of pENSV1 or pENBT1, or with *XbaI/SpeI* and ligation into the *NheI/SpeI* sites of pOSV808 via isocaudomer cloning ^[37]^.

The *vhb, sco691*, accA2BE genes and *ermE*p* promoter were codon-optimized and synthesized as BioBricks® lacking internal *EcoRI, PstI, SpeI*, and *XbaI* restriction sites (Supplementary Information) (Genscript). The *ermE*p* promoter fragment was restriction digested with *EcoRI*/*PstI* and ligated into the *EcoRI*/*PstI* sites of pSB1C3. Expression vectors pENSV1 and pENTG1 were generated as synthetic vectors in a pUC57-mini backbone expressing an origin of transfer sequence (*oriT*), orthogonal actinophage integrase, and drug resistance marker for selection in *Streptomyces coelicolor* M1146 (Supplementary Information). pOSV808 was a gift from Jean-Luc Pernodet (Addgene plasmid # 126601; http://n2t.net/addgene:126601; RRID:Addgene_126601).

### Intergeneric conjugation between E. coli and S. coelicolor

The conjugation donor host *E. coli* ET12567/pUZ8002 was transformed with constructs for mobilization into *Streptomyces coelicolor* M1146::cos16F4iE, as previously described [8]. *Streptomyces coelicolor* recipient strains were grown on SFM agar plates for 5 days to achieve sporulation. In brief, *E. coli* ET12567/pUZ8002 derivative strains harboring expression constructs for conjugation were grown overnight at 37°C in 3 mL of LB liquid media in an orbital shaker. The cultures were centrifuged at 4000 x g for 10 minutes and resuspended in 2 mL of sterile LB media to remove antibiotics. This procedure was repeated twice. In parallel, 3 mL of sterile TSB was added to one plate of well-sporulated *S. coelicolor*, and the spores were gently rubbed off the plate with a sterile spreader and collected in a sterile 15 mL conical centrifuge tube. The spores were heat-shocked at 50°C for 10 minutes and recovered on ice for 10 minutes. 100 µL of spores were mixed with 100 µL of *E. coli* ET12567/pUZ8002 donor cells on SFM media, plated with a sterile spreader, and allowed to dry in a laminar flow hood. The conjugal matings were then incubated at 30°C for 16-20 hours before flooding with 1.0 mL sterile ddH_2_O, nalidixic acid (35 µg mL^-1^), and the appropriate antibiotic(s) for selection of *S. coelicolor* exconjugants. For each transformation, 9 to 12 independent exconjugants were plated to DNA plates supplemented with antibiotics and grown for 4 to 5 days until the formation of vegetative mycelium.

### Production of tetracenomycins and HPLC-MS analysis

For tetracenomycin production experiments, 9 to 12 recombinant *S. coelicolor* exconjugants were plated on ISP4 plates with appropriate antibiotics for 4 to 5 days until the formation of vegetative mycelium. For seed culture fermentations, sterile 15 mL culture tubes (Fisher Scientific) were filled with 2 mL TSB liquid media and inoculated with freshly grown spores (10% v/v), and grown for 2 days. For time-course experiments, sterile 250 mL Erlenmeyer flasks were filled with 25 mL SG-TES liquid media and sterile glass beads (3 mm, 10-20 beads per flask) to inhibit mycelial aggregation. The shake flask fermentations were inoculated with 1 mL seed culture (4% v/v) and grown in an orbital shaker for 5 days. Biomass measurements were recorded using the BugLabs BEH100-Handheld OD scanner, as previously described ^[38]^. The BugLabs BEH100-Handheld OD scanner is a non-invasive optical sensor and emits signals at 850 nm to detect light reflected from cells in the vessel. Experimental determinations were determined based on data obtained from 4 – 6 replicates grown on different days. Data were plotted in figures and independent T-tests were carried out in the GraphPad Prism® 9 Software suite (GraphPad Software, San Diego, CA). 25 mL of cell culture was extracted 1:1 with 25 mL of 0.1% formic acid: ethyl acetate and the organic phase was dried down, resuspended in 4 mL of methanol, and filtered through a 0.45 µm nylon syringe-driven filter.

Analyses and quantification of **1 – 4** were carried out on an Agilent 1260 Infinity II LC/MSD iQ single quadrupole instrument. In brief, 10 µL of the sample was injected via an autosampler onto the sample loop and was separated on a Poroshell 120 Phenyl-Hexyl Column (ID 2.7 µm, 4.6 mm x 100 mm) and was analyzed in gradients of solvent A (0.1% formic acid in water) and solvent B (0.1% formic acid in acetonitrile). The HPLC program used a constant flow rate of 0.5 mL per minute and the following gradient steps: 0 minutes, 95% solvent A and 5% solvent B; 0 – 10 minutes, 95% solvent A and 5% solvent B to 5% solvent A and 95% solvent B; 10 – 13 minutes, held at 5% solvent A and 95% solvent B; 13.1 minutes, re-equilibrate to 95% solvent A and 5% solvent B; 13.1 – 15.1 minutes, 95% solvent A and 5% solvent B. The diode array detector (DAD) was set to monitor UVvis absorbance at 290 nm and 410 nm (i.e. which is selective for tetracenomycins). The ESI-MS was set to scan from 200 *m/z* – 500 *m/z* fragments in positive and negative ionization modes. Single ion monitoring was set-up in ESI-MS negative ionization mode using the following ions: **1** = [M-H] = 457 *m/z*; **2** = [M-H] = 471 *m/z*; **3** = [M-H] = 485 *m/z*, **4** = [M-H] = 487 *m/z*. All biosynthetic samples were compared to authentic standards of **1 – 4**.

## Results

### Heterologous expression of cos16F4iE in Streptomyces coelicolor M1146

Initially, we sought to generate an improved host for improved production of **1** and **5** analogs for downstream antiproliferative activity and drug metabolism studies. The initial host *Streptomyces lividans* TK24 (cos16F4) was based on a pKC505-based cosmid expression system that resulted in the production of 8-DMTC and tetracenomycin B3 at a yield of approximately 15 – 20 mg/L ^[12]^. We routinely worked with this host to generate novel glycosylated tetracenomycins via co-expression of “deoxysugar plasmids” that could direct the biosynthesis of TDP-deoxysugars for glycosylation onto the 8-DMTC aglycone via ElmGT. In our hands, the *S. lividans* TK24 (cos16F4) expression host would experience segregation of cos16F4 or “deoxysugar plasmids” during scaled-up fermentations to isolate new tetracenomycin analogs ^[17]^.

We obtained the integrating vector cos16F4iE for introduction in the improved heterologous expression host *S. coelicolor* M1146. The vector cos16F4iE features the ϕC31 integrase and *attP* attachment site for recombination into the *attB* site of the *S. coelicolor* chromosome. Integration of this cassette could ensure stable expression of the core 8-DMTC biosynthetic genes and could avoid the instability issues observed previously with *S. lividans* TK24 (cos16F4). The *Streptomyces coelicolor* M1146 expression host has several advantages over *S. lividans* TK24, including deletion of four major biosynthetic gene clusters, resulting in a host with fungible metabolism for heterologous expression of type II polyketide synthase gene clusters ^[23]^. Introduction of cos16F4iE into *S. coelicolor* M1146 via intergeneric conjugation resulted in several apramycin-resistant exconjugants that produced an orange-red pigmented color when plated on SFM agar. 12 independent clones were grown up for 5 days and extracted. The methanolic extracts for all twelve strains indicated significant production of **1** and tetracenomycin B3, as compared to an authentic standard of **1** (t_R_ = 8.76 min) ^[12]^. The yield of **1** from the clones ranged from 100 – 160 mg/L, which is a 5 to 8-fold improvement over the original production host. One high-producing clone was carried forward for further experiments.

### Development of Orthogonal BioBrick® Vectors for Integration in Streptomyces coelicolor

Next, we set out to develop a set of orthogonal BioBrick® [RFC-10] vectors for integration of gene cassettes into the chromosome of *Streptomyces coelicolor* M1146::cos16F4iE. We designed new BioBricks® vectors based on the SV1 and TG1 actinophage integrases that could be used for the expression of gene circuits from orthogonal promoters (Supplementary Figure 1). pENSV1 incorporates the SV1 actinophage integrase, the *attP* site, *oriT* for mobilization from *E. coli* ET12567/pUZ8002 via conjugation, and the *aadA* spectinomycin resistance gene for site-specific recombination into the chromosome.

Simultaneously, pENTG1 incorporates the TG1 actinophage integrase, *oriT, attP* site, and the *vph* viomycin resistance gene for single-copy chromosomal engineering (Supplementary Figure 1). In addition, we obtained the BioBrick-compatible vector pOSV808, which includes the VWB actinophage integrase, *attP* site, *oriT*, and the *amilCP* gene for screening of recombinant clones^[37]^.

pENSV1, pENTG1, and pOSV808 were successfully transformed into *S. coelicolor* M1146 and *S. coelicolor* M1146::cos16F4iE via intergeneric conjugation. This demonstrated that these vectors could potentially be useful for shuttling gene cassettes into *S. coelicolor* for pathway engineering. We next used these vectors to clone in different operons for substrate precursor engineering of **1**.

### Engineering precursor metabolite pools to increase production titers of 1

Next, we decided to use the pENSV1, pENTG1, and pOSV808 expression vectors to engineer precursor substrate pools within *S. coelicolor* M1146::cos16F4iE to produce higher levels of **1**. Substrate precursor engineering has been used to increase the production of a variety of aromatic polyketides, including mithramycin, tetracenomycin C, actinorhodin, nogalamycin, and steffimycin ^[39,40]^. The proposed biosynthesis of **1** requires condensation of 1 molecule of acetyl-CoA and 9 molecules of malonyl-CoA via the ElmKLM minimal PKS (Figure 2). Cyclases ElmNI and ElmJ generate the tricyclic tetracenomycin F2, which is cyclized by ElmI to form tetracenomycin F1, oxidatively modified by ElmH to form tetracenomycin D3, and undergoes consecutive *O*-methylations at the 3-*O-*position by ElmNII and at the 9-*O*-position by ElmP ^[41]^. ElmG carries out a triple hydroxylation of the penultimate intermediate to form **1** ^[10]^.

We engineered three different gene cassettes to enhance substrate precursor pools for **1**. First, we engineered the *Streptomyces coelicolor* M145 acetyl-CoA carboxylase complex (i.e. *accA2BE*) under the control of the constitutive *ermE*p* promoter to enhance condensation of acetyl-CoA to malonyl-CoA (Figure 2). This strategy has been successfully used to enhance the production of actinorhodin by 6-fold ^[30]^. Second, we engineered the acyltransferase *sco6196* under the control of the constitutive *ermE*p* promoter to increase carbon flux from triacylglycerols to beta-oxidation, which increases acetyl-CoA precursor supply ^[31]^. polyketide biosynthesis when it is most active. *Sco6196* is a highly active acyltransferase that plays a major role as a “metabolic switch” during stationary phase, which mobilizes triacylglycerols to the beta-oxidation machinery to produce acetyl-CoA, which is then diverted towards polyketide biosynthesis ^[31]^ Lastly, we decided to engineer the *vhb* hemoglobin gene from the obligate aerobe *Vitreoscilla stercoraria* under the control of its oxygen-sensitive promoter. Expression of *vhb* in *S. coelicolor* M1146::cos16F4iE is expected to enhance biomass formation and availability of oxygen for the electron transport chain.

We also hypothesized that the expression of different gene cassettes from unique loci in the *S. coelicolor* chromosome might lead to the identification of “chromosomal position effects”, due to some regions of the chromosome being transcribed more frequently than other regions, which could lead to improved product formation. This strategy was exploited by Bilyk et al. to array production of aranciamycin over an 8-fold range dependent on the *attB* site of recombination ^[42]^. We cloned *vhb, ermE*p-accA2BE*, and *ermE*p-sco6196* onto pENSV1, pENTG1, and pOSV808 to splice the gene constructs into the SV1, TG1, and VWB *attB* sites of *S. coelicolor*. We observed that the engineering the integrating cassettes into the different *attB* sites resulted in decreasing rank order of **1** production titer as follows: SV1 > VWB > TG1. The recombinant strains were grown in SG liquid media in shake flasks for 5 days. After five days, biomass measurements were conducted and the cultures were extracted to determine **1** production titers via HPLC-MS analysis (Figure 3). Each experiment used 4 – 6 biological replicates, which were compared to a standard curve of authentic **1** (Figure 4). The recombinant strains harboring pENSV1-*vhb*, pENSV1-*ermE*p-accA2BE*, and pENSV1-*ermE*p-sco6196* exhibited the highest increases in **1** product titer (Figure 3). The cos16F4iE line produced a mean of 166 mg/L **1**, whereas the cos16F4iE::pENSV1-*vhb* line exhibited 32% increased production of **1** (e.g. 220.3 ± 15.3 mg/L, p = 0.0168), the cos16F4iE::pENSV1-*sco6196* line exhibited 2.2-fold increased production of **1** (e.g. 366.6 ± 67.8 mg/L, p=0.0465), and cos16F4iE::pENSV1-*accA2BE* line exhibited the greatest increase in production titer of **1** of 2.4-fold (e.g. 403 ± 83.6 mg/L, p=0.0304) (Figure 3). HPLC-MS analysis of the different lines revealed an increase in production of **1**, and significant production of the penultimate intermediate tetracenomycin B3 (Figure 4). In addition, the transformation of pOSV808-based constructs resulted in statistically significant increases in titer of **1**. The cos16F4iE::pOSV808-*vhb* strain produced 224.5 ± 18.68 mg/L of **1** (p = 0.0415), cos16F4iE::pOSV808-*sco6196* produced 249.2 ± 11.88 mg/L of **1** (p = 0.0007), and cos16F4iE::pOSV808-*accA2BE* produced 327.3 ± 40.2 mg/L of 1 (p = 0.0131). Surprisingly, engineering of pENTG1-based constructs resulted in statistically significant decreases in **1** production for all combinations attempted. One possible explanation for this could be that recombination of genes into the TG1 *attB* site could be deleterious for the growth of the integrants. TG1 integrates into *sco3658*, which encodes an aminotransferase ^[43]^. In addition, no statistically significant differences were detected in biomass between the control line and the experimental lines, except for a decrease in biomass for the cos16F4iE::pENTG1-*vhb* line (p = 0.0039). This result demonstrates that the engineering of specific genes into different *attB* sites may have unanticipated effects on the growth of the strain, therefore, interrogation of several different *attB* sites might be required to identify the optimal chromosomal locus for expression of a given gene cassette.

**Figure 3.**
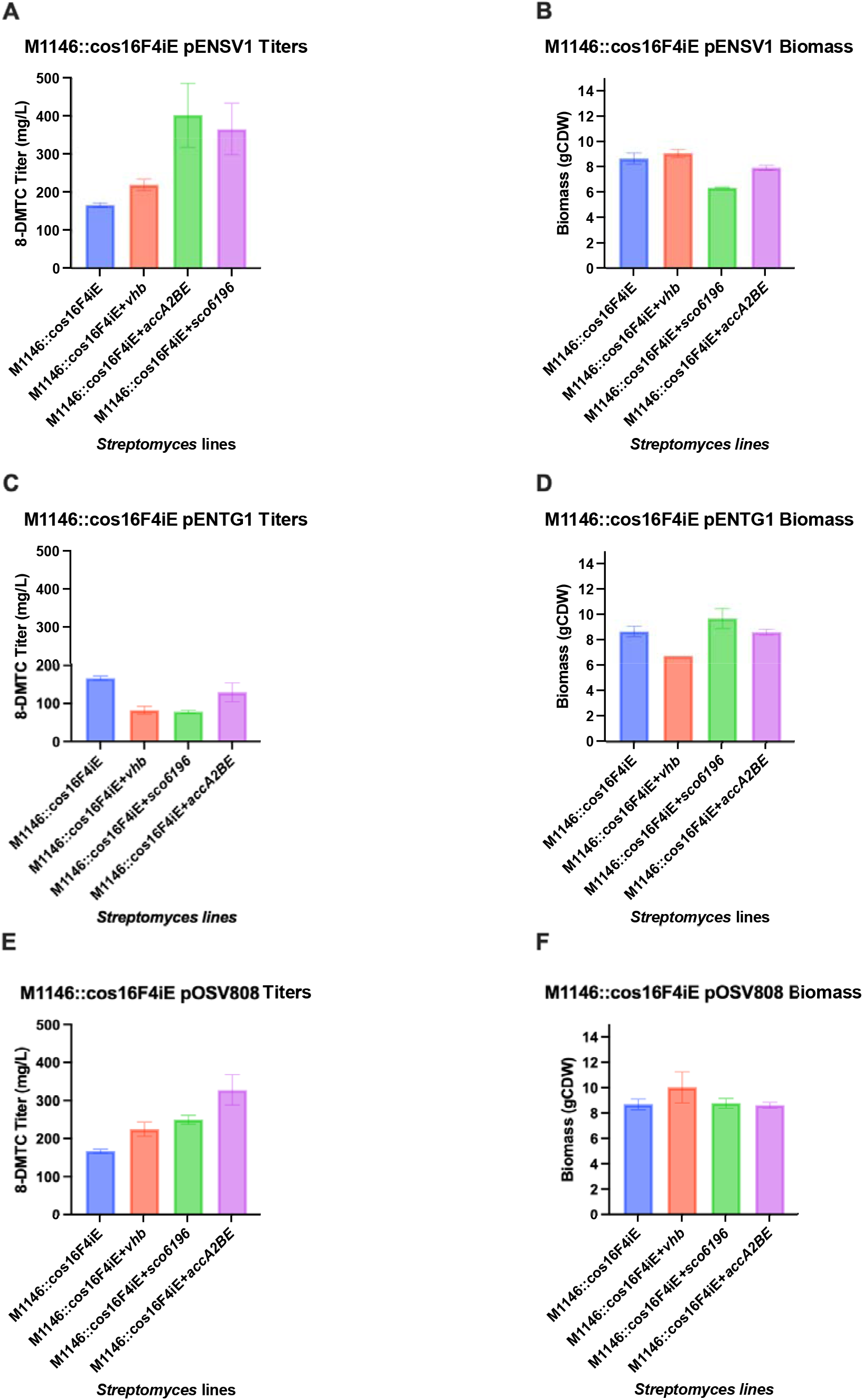
Substrate precursor engineering of *S. coelicolor* M1146::cos16F4iE for production of 1. (A) Production titers of 8-DMTC from lines engineered with pENSV1::*vhb*, pENSV1::*accA2BE*, or pENSV1::*sco6196*. (B) Biomass measurements of lines engineered with pENSV1-based vectors. (C) Production titers of **1** from lines engineered with pENTG1::*vhb*, pENTG1::*accA2BE*, or pENTG1::*sco6196*. (D) Biomass measurements of lines engineered with pENTG1-based vectors. (E) Production titers of **1** from lines engineered with pOSV808::*vhb*, pOSV808::*accA2BE*, or pOSV808::*sco6196*. (F) Biomass measurements of lines engineered with pOSV808-based vectors. Experiments were conducted with 4 – 6 biological replicates. Experimental groups were compared using a t-test to determine statistical significance (*p* < 0.05).

**Figure 4.**
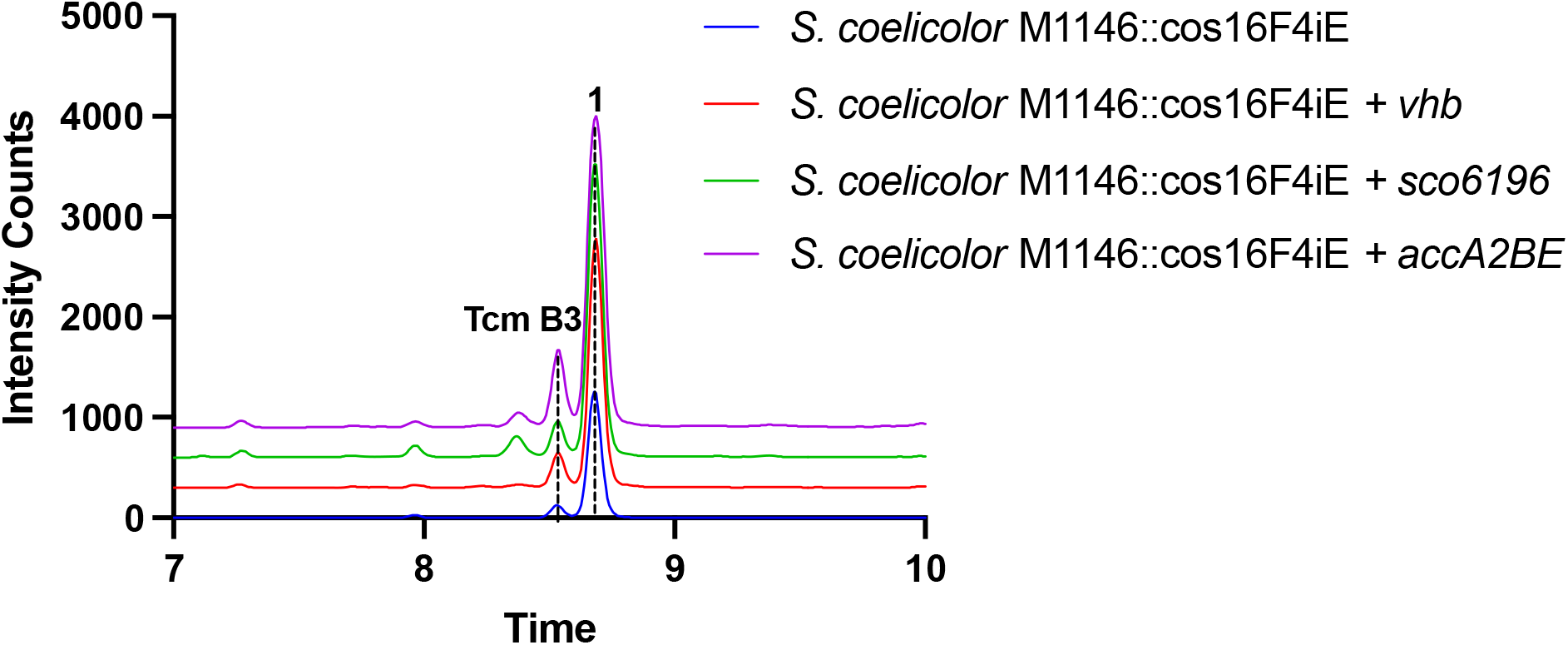
Increased production of 1 via the metabolic engineering of substrate precursor pools. Production of **1** was increased via the expression of a chromosomally-integrated copy of *Vitreoscilla stercoraria* hemoglobin (*vhb*, red trace), *S. coelicolor sco6196* acyltransferase (*sco6196*, green trace), or the *S. coelicolor* acetyl-CoA carboxylase complex (*accA2BE*, purple trace).

### Engineering of tetracenomycin analogs and biosynthesis of tetracenomycin X

We decided to employ our improved production strains for the generation of tetracenomycin analogs **2 - 4** ^[3,7,44]^. The heterologous production of **2** and **3** which feature multiple *O*-methylations, and **4**, which features an additional hydroxyl group, could be useful for downstream structure-activity relationships studies (Figure 1). The biosynthesis for **2** diverges from **1** at tetracenomycin B3: TcmO catalyzes *O*-methylation at the 8-position to form tetracenomycin E, which is *O*-methylated at the 9-position by ElmP (i.e., TcmP homolog from *S. olivaceus* Tü 2353) to afford tetracenomycin A2, which is hydroxylated at 4, 4a, and 12a positions by ElmG (i.e., TcmG homolog from *S. olivaceus* Tü 2353) (Figure 2). **1** biosynthesis does not undergo 8-*O*-methylation (since the 8-position is glycosylated by ElmGT), therefore, tetracenomycin B3 is *O*-methylated at 9-position by ElmP then hydroxylated by ElmG to afford **1**. In summation, **1** is a shunt product with respect to the 8-*O*-methyltransferase TcmO, and the capacity of the elloramycin pathway enzymes to modify late-stage noncanonical tetracenomycin substrates is unknown. In specific, the capability for ElmP to *O*-methylate tetracenomycin E or ElmG to hydroxylate tetracenomycin A2 to **2** is uncertain.

To test this hypothesis, we synthesized a codon-optimized version of the *tcmO* gene from the *S. glaucescens* GLA.0 pathway and expressed it under the control of the *sf14p* promoter in a pENSV1 vector in *S. coelicolor* M1146::cos16F4iE and *S. coelicolor* M1146::cos16F4iE::pOSV808-*accA2BE*. Both strains accumulated minor quantities of **2** as determined via comparison to an authentic standard of **2** (t_R_ = 9.39 min) (Figure 5). We also sought to characterize the recently sequenced *tcmO* homolog from the **3** producer *Amycolatopsis spp*. A23. We synthesized the codon-optimized version of the *tcmO* homolog from *Amycolatopsis spp*. A23 and similarly expressed it under the control of the *sf14p* promoter in a pENSV1 vector in *S. coelicolor* M1146::cos16F4iE and *S. coelicolor* M1146::cos16F4iE ::pOSV808-*accA2BE*. Again, both strains accumulated minor quantities of **2** as expected. This result shows that the *tcmO* homolog from *Amycolatopsis spp*. A23 encodes a tetracenomycin B3 8-*O*-methyltransferase. In addition, this result demonstrates that ElmP is capable of *O*-methylating tetracenomycin E and ElmG is flexible enough to convert tetracenomycin A2 to **2**. It was difficult to determine the production titer for these metabolites since the amount of **2** produced by each strain was <1% of the total amount of TCMs detected in the HPLC chromatogram. We were able to quantify relative production based on filtering the data in single-ion monitoring (SIM) in ESI negative ionization mode by searching for the [M-H] = 471 *m/z* ion.

**Figure 5.**
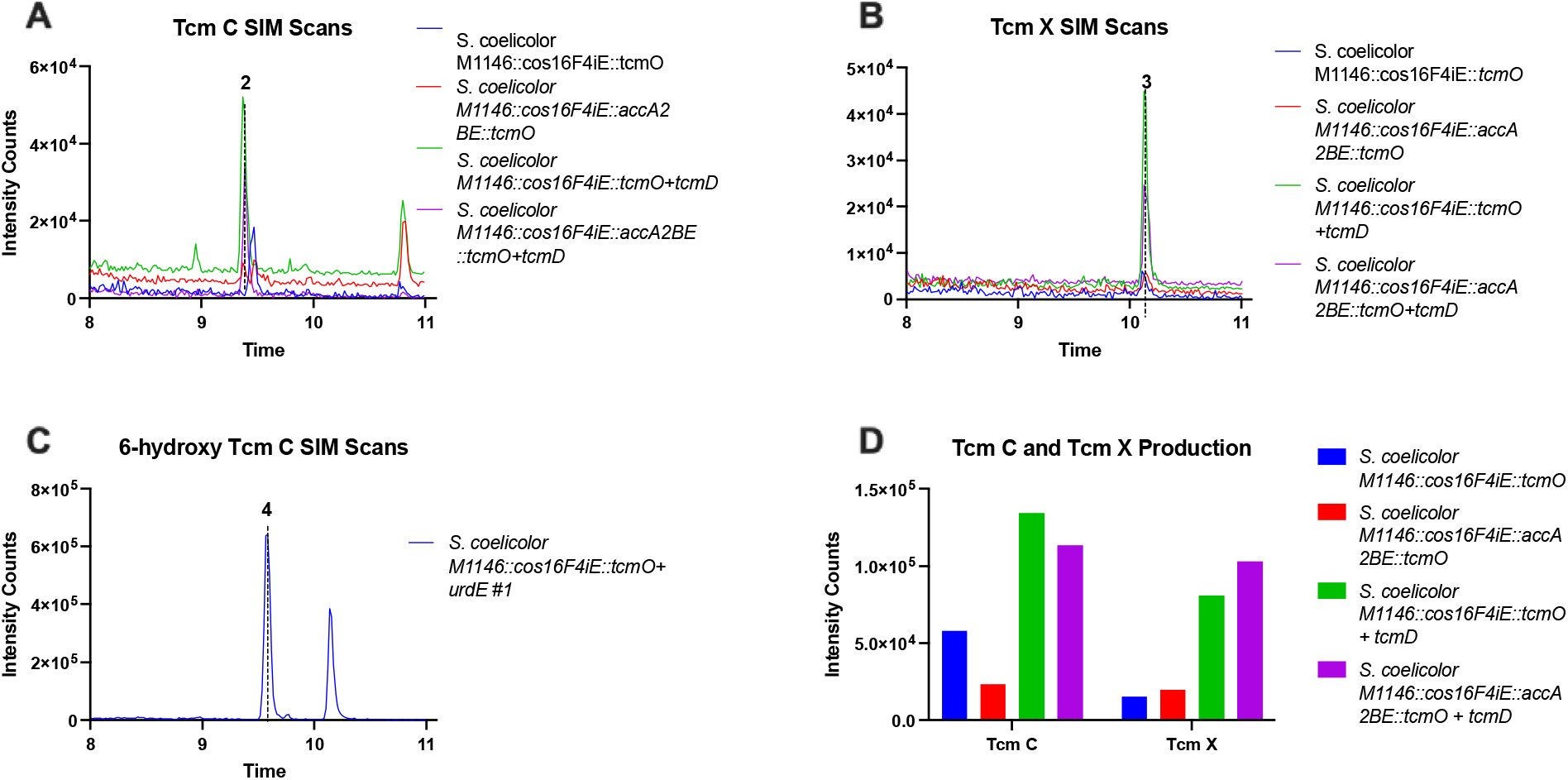
Production of tetracenomycin analogs. (Panel A) Chromatogram traces of lines producing **2** as analyzed in ESI-MS negative ion SIM mode: [M-H] = 471 *m/z*. (Panel B) Chromatogram traces of lines producing **3** as analyzed in ESI-MS negative ion SIM mode: [M-H] = 485 *m/z*. (Panel C) Chromatogram traces of lines producing **4** as analyzed in ESI-MS positive ion SIM mode: [M+H] = 489 *m/z*. (Panel D) Quantification of relative amounts of **2** and **3** from different production lines based on intensity counts in ESI-MS negative ion SIM mode.

Secondly, we decided to build on this previous result by incorporating the *urdE* oxygenase from the urdamycin pathway to hydroxylate **2** to **4** (Figure 2). UrdE was previously shown to accept **2** as an alternative substrate to its preferred angucyclinone substrate and can carry out hydroxylation at the 6-position of **2** ^[44]^. We synthesized a codon-optimized version of *urdE* and cloned it under the *p* promoter into our pENSV1-*sf14p-tcmO* vector and expressed it in *S. coelicolor* M1146::cos16F4iE. Analysis of methanolic extracts from this strain resulted in the detection of **4** in SIM ESI positive ion mode using the [M-H] = 489 *m/z* ion as compared to an authentic standard of **4** (t_R_ = 9.64 min) (Figure 5). The yield of this compound was very low, <1% of total TCMs, most likely owing to the relatively high level of metabolic flux towards **1** production. All attempts to transform the pENSV1-*sf14p-tcmO-sf14p-urdE* construct into *S. coelicolor* M1146::cos16F4iE::pOSV808 resulted in transformants that failed to grow on agar plates. One possible explanation for this observation could be that the ElmE elloramycin permease does not actively transport **4** outside of the cell, which could lead to toxicity due to the intracellular accumulation of **4**.

Thirdly, we decided to investigate the biosynthesis of **3**, which is previously uncharacterized. We hypothesized that an *S*-adenosyl-L-methionine-dependent 12-*O-* methyltransferase (i.e. SAM-dependent *O-*MT) methylates **2** to **3** (Figure 2). Further investigation in SIM ESI negative ion mode of the extracts from *S. coelicolor* M1146::cos16F4iE::pENSV1-*tcmO* revealed the presence of another methylated tetracenomycin with a mass of 486 *amu* and a later elution profile than **2** (t_R_ = 10.15 min) (Figure 4). We identified the peak as **3** as compared to an authentic standard. This result demonstrates that the elloramycin 12-*O*-methyltransferase, ElmD, is capable of methylating **2** to form **3**. Knowing that the **3** gene cluster from *Amycolatopsis* spp. A23 would likely include an ElmD paralogue, we conducted a translated nucleotide basic local alignment search tool (BLASTX®) search of the *Amycolatopsis* spp. A23 genome with the ElmD nucleotide sequence as a search query. The search resulted in the identification of a 296 amino acid SAM-dependent *O*-MT (Accession Number WP_155542896.1) with significant sequence homology (e.g. 54% identical/67% similar) to ElmD. We decided to call this enzyme TcmD, and we proceeded to synthesize a codon-optimized version of *tcmD* for co-expression in our pENSV1-*sf14p-tcmO* construct. We cloned *tcmD* under a copy of an additional *sf14p* promoter and spliced it at the 3’-end of *tcmO*. We expressed the resulting pENSV1-*sf14p-tcmO-sf14p-tcmD* construct in both *S. coelicolor* M1146::cos16F4iE and *S. coelicolor* M1146::cos16F4iE::pOSV808-*accA2BE* (Figure 5). While quantification of the production titer of **3** was not possible due to the low level of production, <1% of all TCMs, we were able to determine relative production amounts of **2** and **3** via intensity counts from the mass spectrometer. Expression of *tcmO* itself lead to approximately equimolar production of **2** and **3** in extracts from *S. coelicolor* M1146::cos16F4iE ::pOSV808-*accA2BE*::pENSV1-*sf14p-tcm*O. Co-expression of *tcmD* resulted in a ten-fold increase in **3** production titer, as well as a significant increase in **2** production. The highest yields resulted from co-expression of *tcmO* and *tcmD* in the line harboring the acetyl-CoA carboxylase complex, which highlights the fact that the increased production of **2** and **3** was due to the increased substrate precursor pools and resultant metabolic flux through the engineered tetracenomycin pathway in this strain. In summation, to the best of our knowledge, this is the first report in which *tcmD* has been characterized via heterologous expression as the 12-*O*-MT responsible for the biosynthesis of **3**.

## Discussion

In this report, we developed a series of orthogonal integrating vectors based on the SV1, TG1, and VWB actinophage integrases and used these vectors to engineer in *vhb* hemoglobin, *accA2BE* acetyl-CoA carboxylase, and *sco6196* acyltransferase gene cassettes. This multiplexed metabolic engineering strategy resulted in improved production strains of *S. coelicolor* M1146::cos16F4iE, especially those lines expressing *accA2BE* or *sco6196*, which resulted in the highest production titer of 486 mg/L. Previously, Li et al. engineered the **2** biosynthetic pathway in a knockout mutant of the industrial monensin producer, *Streptomyces cinnomonaeus* ^[45]^. The highest reported production of **3** in this strain was 440 mg/L, which indicates that our production methodology compares favorably with this industrial host. Furthermore, industrial hosts often result from iterative cycles of random mutagenesis and screening for mutants with desired production characteristics. Our methodology provides a rational approach for improving type II polyketide production titers based on several complementary approaches, including overexpression of the acetyl-CoA carboxylase complex to enhance malonyl-CoA concentration, overexpression of *sco6196* to enhance acetyl-CoA levels, and overexpression of the *vhb* hemoglobin to enhance oxygen concentrations in submerged liquid fermentation, which could boost cellular metabolism, as well as enhance biosynthetic oxygenation steps.

We used this enhanced production platform as a showcase for combinatorial biosynthesis of tetracenomycin analogs **2 – 4**. These analogs are thought to be more valuable anticancer compounds than **1**, owing to the *O*-methyl groups at 8- and 12-positions, which enhance binding to the large ribosomal polypeptide exit channel ^[3,5]^. Engineering of *tcmO* orthologs from two different actinomycetes, *S. glaucescens* GLA.0 and *Amycolatopsis* sp. A23 resulted in the production of **2** and **3**. The heterologous expression of *urdE* also resulted in the production of **4**, as previously described ^[44]^. Most importantly, the heterologous expression of the newly characterized *tcmD* gene resulted in a ten-fold increase in production of **3**, which provides good evidence for its role as a tetracenomycin C 12-*O*-methyltransferase in the **3** biosynthetic pathway. The utility of this production method is diminished, however, by the significant metabolic flux away from **2 – 4** production to production of **1** at the tetracenomycin B3 step. Tetracenomycin B3 likely represents a branch point for the glycosylated elloramycins and methylated tetracenomycins. Future studies should focus on engineering combinations of *tcmD* and *urdE* in the *Streptomyces glaucescens* GLA.0 tetracenomycin C wildtype producer. In this strain, we expect that higher production titers of **3** and **4** could be realized, in addition to the potential for producing new tetracenomycin analogs.

## Acknowledgments

Research reported in this publication was supported by theNational Cancer Institute of the National Institutes of Health under Award No. R15CA252830 (to S.E.N.) and the National Science Foundation under Grant No. ENG-2015951 (to S.E.N.). The authors also acknowledge the Ferris State University College of Pharmacy and the College of Pharmacy Alumni Board for financial support for the purchase of the Agilent 1260 Infinity II iQ HPLC-MS instrument used in this study. The authors also thank Dr. Jose Salas (University of Oviedo, Spain) for cosmid cos16F4iE. The authors also acknowledge Dr. Juan Pablo-Escribano and Dr. Mark Buttner (John Innes Center, Norwich, United Kingdom) for the provision of *Streptomyces coelicolor* M1146. The authors also thank Dr. Khaled A. Shaaban of the Center for Pharmaceutical Research and Innovation, University of Kentucky College of Pharmacy for providing chemical standards for **1** and **4**. The authors also acknowledge the National Cancer Institute Developmental Therapeutics Program for providing chemical standards for **2** and **3**.

S.E.N. conceptualized the study, acquired funding, supervised all aspects of the study, and wrote the manuscript. K.V.B., J.T.N., N.M.G., and K.K.R. carried out experiments (equal). All authors reviewed the data and edited the manuscript.

## Conflict of Interest

Material published in this report is covered under U.S. Patent Application No. 16/015,821 to Ferris State University.

## References

1. Egert, E., Noltemeyer, M., Siebers, J., Rohr, J., & Zeeck, A. (2012). The structure of tetracenomycin C. The Journal of Antibiotics, 45(7), 1190–1192. https://doi.org/10.7164/antibiotics.45.1190

2. Zeeck, A., Reuschenbach, P., ZäHner, H., & Rohr, J. (1985). Metabolic products of microorganisms. 2251) elloramycin, a new anthracycline-like antibiotic from streptomyces olivaceus isolation, characterization, structure and biological properties. The Journal of Antibiotics, 38(10), 1291–1301. https://doi.org/10.7164/antibiotics.38.1291

3. Rohr, J., & Zeeck, A. (1990). Structure-activity relationships of elloramycin and tetracenomycin C. The Journal of Antibiotics, 43(9), 1169–1178. https://doi.org/10.7164/antibiotics.43.1169

4. Qiao, X., Gan, M., Wang, C., Liu, B., Shang, Y., Li, Y., & Chen, S. (2019). Tetracenomycin X Exerts Antitumour Activity in Lung Cancer Cells through the Downregulation of Cyclin D1. Marine Drugs, 17(1), 63. https://doi.org/10.3390/md17010063

5. Osterman, I. A., Wieland, M., Maviza, T. P., Lashkevich, K. A., Lukianov, D. A., Komarova, E. S., Zakalyukina, Y. v., Buschauer, R., Shiriaev, D. I., Leyn, S. A., Zlamal, J. E., Biryukov, M. v., Skvortsov, D. A., Tashlitsky, V. N., Polshakov, V. I., Cheng, J., Polikanov, Y. S., Bogdanov, A. A., Osterman, A. L., … Sergiev, P. v. (2020). Tetracenomycin X inhibits translation by binding within the ribosomal exit tunnel. Nature Chemical Biology, 1–7. https://doi.org/10.1038/s41589-020-0578-x

6. Ortseifen, V., Kalinowski, J., Pühler, A., & Rückert, C. (2017). The complete genome sequence of the actinobacterium Streptomyces glaucescens GLA.O (DSM 40922) carrying gene clusters for the biosynthesis of tetracenomycin C, 5’-hydroxy streptomycin, and acarbose. Journal of Biotechnology, 262, 84–88. https://doi.org/10.1016/j.jbiotec.2017.09.008

7. Motamedi, H., & Hutchinsont, C. R. (1987). Cloning and heterologous expression of a gene cluster for the biosynthesis of tetracenomycin C, the anthracycline antitumor antibiotic of Streptomyces glaucescens (antibiotic resistance/pigment production genes). In Biochemistry (Vol. 84).

8. Thompson, T. B., Katayama, K., Watanabe, K., Hutchinson, C. R., & Rayment, I. (2004). Structural and functional analysis of tetracenomycin F2 cyclase from Streptomyces glaucescens: A type II polyketide cyclase. Journal of Biological Chemistry, 279(36), 37956– 37963. https://doi.org/10.1074/jbc.M406144200

9. Ames, B. D., Korman, T. P., Zhang, W., Smith, P., Vu, T., Tang, Y., & Tsai, S. C. (2008). Crystal structure and functional analysis of tetracenomycin ARO/CYC: Implications for cyclization specificity of aromatic polyketides. Proceedings of the National Academy of Sciences of the United States of America, 105(14), 5349–5354. https://doi.org/10.1073/pnas.0709223105

10. Shen, B., & R, H. (1994). Triple Hydroxylation of Tetracenomycin A2 to tetraenomycin C in Streptomyces glaucescens. The Journal of Biological Chemistry, 1326(25), 30726–30733. http://www.jbc.org/content/269/48/30726.full.pdf

11. Yue, S., Motamedi, H., Wendt-Pienkowski, E., & Hutchinson, C. R. (1986). Anthracycline Metabolites of Tetracenomycin C-Nonproducing Streptomyces glaucescens Mutants. In JOURNAL OF BACTERIOLOGY. http://jb.asm.org/

12. Decker, H., Rohr, J., Motamedi, H., Zähner, H., & Hutchinson, C. R. R. (1995). Identification of Streptomyces olivaceus Tü 2353 genes involved in the production of the polyketide elloramycin. In Gene (Vol. 166, Issue 1). Elsevier. https://doi.org/10.1016/0378-1119(95)00573-7

13. Blanco, G., Patallo, E. P., Braña, A. F., Trefzer, A., Bechthold, A., Rohr, J., Méndez, C., & Salas, J. A. (2001). Identification of a sugar flexible glycosyltransferase from Streptomyces olivaceus, the producer of the antitumor polyketide elloramycin. Chemistry & Biology, 8(3), 253–263. https://doi.org/10.1016/S1074-5521(01)00010-2

14. Rodriguez, L., Oelkers, C., Aguirrezabalaga, I., Braña, A. F., Rohr, J. J., Méndez, C., Salas, J. A., Brana, a F., Rohr, J. J., Mendez, C., & Salas, J. A. (2000). Generation of hybrid elloramycin analogs by combinatorial biosynthesis using genes from anthracycline-type and macrolide biosynthetic pathways. Journal of Molecular Microbiology and Biotechnology, 2(3), 271– 276. http://www.ncbi.nlm.nih.gov/pubmed/10937435

15. Rodríguez, L., Aguirrezabalaga, I., Allende, N., Braña, A. F., Méndez, C., & Salas, J. A. (2002). Engineering Deoxysugar Biosynthetic Pathways from Antibiotic-Producing Microorganisms. Chemistry & Biology, 9(6), 721–729. https://doi.org/10.1016/s1074-5521(02)00154-0

16. Pérez, M., Lombó, F., Baig, I., Braña, A. F., Rohr, J., Salas, J. A., & Méndez, C. (2006). Combinatorial biosynthesis of antitumor deoxysugar pathways in Streptomyces griseus: Reconstitution of “unnatural natural gene clusters” for the biosynthesis of four 2,6-D-dideoxyhexoses. Applied and Environmental Microbiology, 72(10), 6644–6652. https://doi.org/10.1128/AEM.01266-06

17. Eric Nybo, S., Shabaan, K. A., Kharel, M. K., Sutardjo, H., Salas, J. A., Méndez, C., & Rohr, J. (2012). Ketoolivosyl-tetracenomycin C: A new ketosugar bearing tetracenomycin reveals new insight into the substrate flexibility of glycosyltransferase ElmGT. Bioorganic and Medicinal Chemistry Letters, 22(6), 2247–2250. https://doi.org/10.1016/j.bmcl.2012.01.094

18. Pérez, M., Lombó, F., Zhu, L., Gibson, M., Braña, A. F., Rohr, J., Salas, J. A., & Méndez, C. (2005). Combining sugar biosynthesis genes for the generation of <scp>l</scp> - and <scp>d</scp>-amicetose and formation of two novel antitumor tetracenomycins. Chem. Commun., 0(12), 1604–1606. https://doi.org/10.1039/B417815G

19. Fischer, C., Rodríguez, L., Patallo, E. P., Lipata, F., Braña, A. F., Méndez, C., Salas, J. A., & Rohr, J. (2002). Digitoxosyltetracenomycin C and glucosyltetracenomycin C, two novel elloramycin analogues obtained by exploring the sugar donor substrate specificity of glycosyltransferase ElmGT. Journal of Natural Products, 65(11), 1685–1689. https://doi.org/10.1021/np020112z

20. Lombó, F., Gibson, M., Greenwell, L., Braña, A. F., Rohr, J., Salas, J. A., & Méndez, C. (2004). Engineering biosynthetic pathways for deoxysugars: branched-chain sugar pathways and derivatives from the antitumor tetracenomycin. Chemistry & Biology, 11(12), 1709–1718. https://doi.org/10.1016/j.chembiol.2004.10.007

21. Birch, A., Hausler, A., & Hutter, R. (1990). Genome Rearrangement and Genetic Instability in Streptomyces spp (Vol. 172, Issue 8). http://jb.asm.org/

22. Kormanec, J., Rezuchova, B., Homerova, D., Csolleiova, D., Sevcikova, B., Novakova, R., & Feckova, L. (2019). Recent achievements in the generation of stable genome alterations/mutations in species of the genus Streptomyces. In Applied Microbiology and Biotechnology (Vol. 103, Issue 14, pp. 5463–5482). Springer Verlag. https://doi.org/10.1007/s00253-019-09901-0

23. Gomez-Escribano, J. P., & Bibb, M. J. (2011). Engineering Streptomyces coelicolor for heterologous expression of secondary metabolite gene clusters. Microbial Biotechnology, 4(2), 207–215. https://doi.org/10.1111/j.1751-7915.2010.00219.x

24. Gregory, M. A., Till, R., & Smith, M. C. M. M. (2003). Integration site for Streptomyces phage φBT1 and development of site-specific integrating vectors. Journal of Bacteriology, 185(17), 5320–5323. https://doi.org/10.1128/JB.185.17.5320-5323.2003

25. Morita, K., Yamamoto, T., Fusada, N., Komatsu, M., Ikeda, H., Hirano, N., & Takahashi, H. (2009). The site-specific recombination system of actinophage TG1. FEMS Microbiology Letters, 297(2), 234–240. https://doi.org/10.1111/j.1574-6968.2009.01683.x

26. van Mellaert, L., Mei, L., Lammertyn, E., Schacht, S., & Anne, J. (1998). Site-specific integration of bacteriophage VWB genome into Streptomyces venezuelae and construction of a VWB-based integrative vector. Microbiology, 144(12), 3351–3358. https://doi.org/10.1099/00221287-144-12-3351

27. Fayed, B., Younger, E., Taylor, G., & Smith, M. C. M. (2014). A novel Streptomyces spp. integration vector derived from the S. venezuelae phage, SV1. BMC Biotechnology, 14(1), 51. https://doi.org/10.1186/1472-6750-14-51

28. Khosla, C., & Bailey, J. E. (1988). The Vitreoscilla hemoglobin gene: Molecular cloning, nucleotide sequence and genetic expression in Escherichia coli. MGG Molecular & General Genetics, 214(1), 158–161. https://doi.org/10.1007/BF00340195

29. Magnolo, S. K., Leenutaphong, D. L., Demodena, J. A., Curtis, J. E., Bailey, J. E., Galazzo, J. L., & Hughes, D. E. (1991). Actinorhodin production by streptomyces coelicolor and growth of streptomyces lividans are improved by the expression of a bacterial hemoglobin. Bio/Technology, 9(5), 473–476. https://doi.org/10.1038/nbt0591-473

30. Ryu, Y. G., Butler, M. J., Chater, K. F., & Lee, K. J. (2006). Engineering of primary carbohydrate metabolism for increased production of actinorhodin in Streptomyces codicolor. Applied and Environmental Microbiology, 72(11), 7132–7139. https://doi.org/10.1128/AEM.01308-06

31. Wang, W., Li, S., Li, Z., Zhang, J., Fan, K., Tan, G., Ai, G., Lam, S. M., Shui, G., Yang, Z., Lu, H., Jin, P., Li, Y., Chen, X., Xia, X., Liu, X., Dannelly, H. K., Yang, C., Yang, Y., … Zhang, L. (2020). Harnessing the intracellular triacylglycerols for titer improvement of polyketides in Streptomyces. Nature Biotechnology, 38(1), 76–83. https://doi.org/10.1038/s41587-019-0335-4

32. Sambrook, J., & W Russell, D. (2001). Molecular Cloning: A Laboratory Manual. Cold Spring Harbor Laboratory Press, Cold Spring Harbor, NY, 999. http://books.google.com/books?id=YTxKwWUiBeUC&printsec=frontcover%5Cnpapers2://publication/uuid/BBBF5563-6091-40C6-8B14-06ACC3392EBB

33. MacNeil, D. J., Gewain, K. M., Ruby, C. L., Dezeny, G., Gibbons, P. H., & MacNeil, T. (1992). Analysis of Streptomyces avermitilis genes required for avermectin biosynthesis utilizing a novel integration vector. Gene, 111(1), 61–68. https://doi.org/10.1016/0378-1119(92)90603-M

34. Kieser, T., Bibb, M. J., Buttner, M. J., Chater, K. F., & Hopwood, D. A. (2000). Practical Streptomyces Genetics. In John Innes Centre Ltd. (p. 529). https://doi.org/10.4016/28481.01

35. Knight, T. (2003). Idempotent Vector Design for Standard Assembly of Biobricks Idempotent Vector Design for Standard Assembly of Biobricks. Structure, 1–11. https://doi.org/http://hdl.handle.net/1721.1/21168

36. Shetty, R. P., Endy, D., & Knight, T. F. (2008). Engineering BioBrick vectors from BioBrick parts. Journal of Biological Engineering, 2(1), 5. https://doi.org/10.1186/1754-1611-2-5

37. Aubry, C., Pernodet, J. L., & Lautru, S. (2019). Modular and integrative vectors for synthetic biology applications in Streptomyces spp. Applied and Environmental Microbiology, 85(16). https://doi.org/10.1128/AEM.00485-19

38. Nakouti, I., & Hobbs, G. (2015). The Application of an On-Line Optical Sensor to Measure Biomass of a Filamentous Bioprocess. Fermentation, 1(1), 79–85. https://doi.org/10.3390/fermentation1010079

39. Zabala, D., Braña, A. F., Salas, J. A., & Méndez, C. (2016). Increasing antibiotic production yields by favoring the biosynthesis of precursor metabolites glucose-1-phosphate and/or malonyl-CoA in Streptomyces producer strains. The Journal of Antibiotics, 69(3), 179–182. https://doi.org/10.1038/ja.2015.104

40. Zabala, D., Braña, A. F., Flórez, A. B., Salas, J. A., & Méndez, C. (2013). Engineering precursor metabolite pools for increasing production of antitumor mithramycins in Streptomyces argillaceus. Metabolic Engineering, 20, 187–197. https://doi.org/10.1016/j.ymben.2013.10.002

41. Ramos, A., Lombo, F., Brana, A. F., Rohr, J., Mendez, C., & Salas, J. A. (2008). Biosynthesis of elloramycin in Streptomyces olivaceus requires glycosylation by enzymes encoded outside the aglycon cluster. Microbiology, 154(3), 781–788. https://doi.org/10.1099/mic.0.2007/014035-0

42. Bilyk, B., Horbal, L., & Luzhetskyy, A. (2017). Chromosomal position effect influences the heterologous expression of genes and biosynthetic gene clusters in Streptomyces albus J1074. Microbial Cell Factories, 16(1), 5. https://doi.org/10.1186/s12934-016-0619-z

43. Myronovskyi, M., & Luzhetskyy, A. (2013). Genome engineering in actinomycetes using site-specific recombinases. In Applied Microbiology and Biotechnology (Vol. 97, Issue 11, pp. 4701–4712). https://doi.org/10.1007/s00253-013-4866-1

44. Decker, H., & Haag, S. (1995). Cloning and characterization of a polyketide synthase gene from Streptomyces fradiae Tü2717, which carries the genes for biosynthesis of the angucycline antibiotic urdamycin A and a gene probably involved in its oxygenation. Journal of Bacteriology, 177(21), 6126–6136. https://doi.org/10.1128/jb.177.21.6126-6136.1995

45. Li, C., Hazzard, C., Florova, G., & Reynolds, K. A. (2009). High titer production of tetracenomycins by heterologous expression of the pathway in a Streptomyces cinnamonensis industrial monensin producer strain. Metabolic Engineering, 11(6), 319– 327. https://doi.org/10.1016/j.ymben.2009.06.004

