## Supplementary Information for "A BioBricks^®^ toolbox for multiplexed metabolic engineering of central carbon metabolism in the tetracenomycin pathway"

**Supplementary Table 1 Strains and plasmids used in this study.**

| Strain or plasmid | Genotype and relevant characteristics | Reference/Accession No. |
| --- | --- | --- |
| Plasmids |  |  |
| pSET152 | *aac3(IV)*, *oriT*, *lacZα*, ΦC31int, *attP*, MCS | (Bierman et al., 1992) |
| pSET152-BBa | BioBricks®-compatible vector; *aac3(IV)^R^*, *oriT*, φC31int, *attP*, MCS | This study. |
| pENSV1 | BioBricks®-compatible vector; *aadA^R^*, *bla^R^*, *oriT*, φSV1int, *attP* | This study. Accession No. MZ547130 |
| pENTG1 | BioBricks®-compatible vector; *vph^R^*, *bla^R^*, *oriT*, φTG1int, *attP* | This study. Accession No. MZ547131 |
| pOSV808 | BioBricks®-compatible vector; *hph^R^,* *oriT*, VWBint, *attP*, *amilCFP* | (Aubry et al., 2019) |
| pUC57-*vhb* | BioBrick® codon-optimized hemoglobin gene from *Vitreoscilla stercoraria* in pUC57, Amp^R^. | GenScript. Accession No. MZ547132 |
| pUC57-*accA2BE* | BioBrick® codon-optimized acetyl-CoA carboxylase complex genes from *Streptomyces coelicolor* in pUC57, Amp^R^. | GenScript. Accession No. MZ547133 |
| pUC57-*sco6196* | BioBrick® codon-optimized *sco6196* acyltransferase from *Streptomyces coelicolor* in pUC57, Amp^R^. | GenScript. Accession No. MZ547134 |
| pUC57-*tcmO* | BioBrick® codon-optimized *tcmO* gene from *Streptomyces glaucescens* GLA.0 in pUC57, Amp^R^. | GenScript. Accession No. MZ567174 |
| pUC57-*tcmO* | BioBrick® codon-optimized *tcmO* gene from *Amycolatopsis sp.* A23 in pUC57, Amp^R^. | GenScript. Accession No. MZ567173 |
| pUC57-*tcmD* | BioBrick® codon-optimized *tcmD* gene from *Amycolatopsis sp.* A23 in pUC57, Amp^R^. | GenScript. Accession No. MZ567175 |
| pUC57-*urdE* | BioBrick® codon-optimized *urdE* gene from *Streptomyces fradiae* Tü 2717 | Genscript.  Accession No. MZ567172 |
| pSV1-*vhb* | *vhb* fragment cloned into pENSV1 | This study. |
| pTG1-*vhb* | *vhb* fragment cloned into pENTG1 | This study. |
| pOSV808-*vhb* | *vhb* fragment cloned into pOSV808 | This study. |
| pSV1-*accA2BE* | *ermE**p-*accA2BE* fragment cloned into pENSV1 | This study. |
| pTG1-*accA2BE* | *ermE**p-*accA2BE* fragment cloned into pENTG1 | This study. |
| pOSV808-*accA2BE* | *ermE**p-*accA2BE* fragment cloned into pOSV808 | This study. |
| pSV1-*sco6196* | *ermE**p-*sco6196* fragment cloned into pENSV1 | This study. |
| pTG1-*sco6196* | *ermE**p-*sco6196* fragment cloned into pENTG1 | This study. |
| pOSV808-*sco6196* | *ermE**p-*sco6196* fragment cloned into pOSV808 | This study. |
| *E. coli* strains |  |  |
| *E. coli* JM109 | endA1 glnV44 thi-1 relA1 gyrA96 recA1 mcrB+Δ(lac-proAB) e14-[F' traD36 proAB+ lacIq lacZ ΔM15] sdR17(rK-mK+); general cloning host | (Yanisch-Perron et al., n.d.) |
| *E. coli* ET12567 | *F- dam*13::Tn9 *dcm*6 *hsdM* *hsdR* *zjj-*202::Tn10 *recF*143 *galK2* *galT22* *ara*-14 *lac*Y1 *xyl*-5 *leuB6* *thi-*1 *tonA*31 *rpsL*136 *his*G4 *tsx-*78 *mtl*-1 *glnV*44 | (Flett et al., 1997) |
| *E. coli* ET12567/ (pUZ8002) | *F- dam*13::Tn9 *dcm*6 *hsdM* *hsdR* *zjj-*202::Tn10 *recF*143 *galK2* *galT22* *ara*-14 *lac*Y1 *xyl*-5 *leuB6* *thi-*1 *tonA*31 *rpsL*136 *his*G4 *tsx-*78 *mtl*-1 *glnV*44; *tra, neo^R^, RP4*; *E. coli*-*Streptomyces* intergeneric conjugation host | (Flett et al., 1997) |
| *Streptomyces strains* |  |  |
| *S. coelicolor* M1146 | SCP1- SCP2-; wildtype actinorhodin producer | (Bentley et al., 2002) |
| *S. coelicolor* M1146::cos16F4iE | *S. coelicolor* M1146 lysogenized with cos16F4iE. Produces 8-DMTC. | This study. |

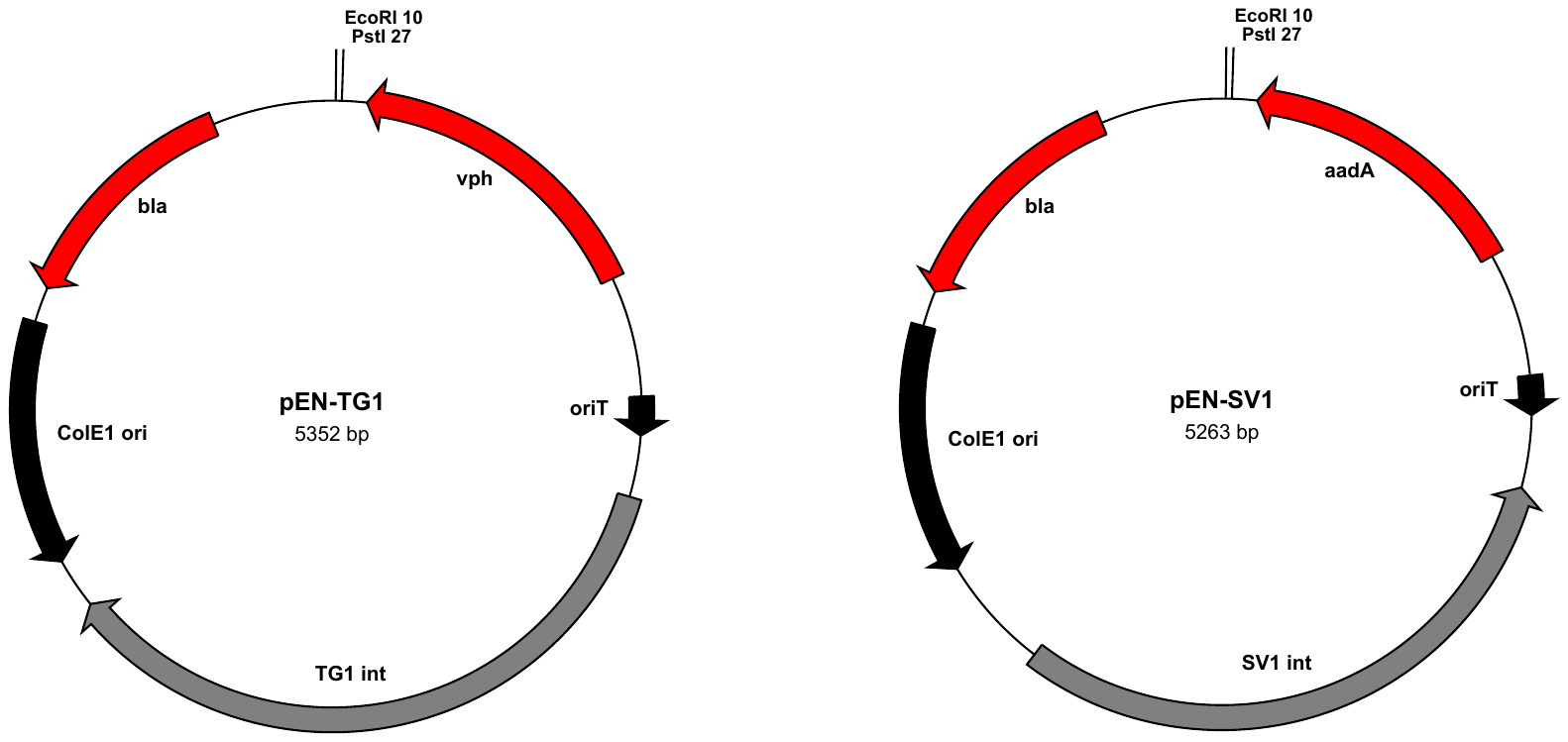

**Supplementary Figure 1** Plasmid maps for pENTG1 and pENSV1.

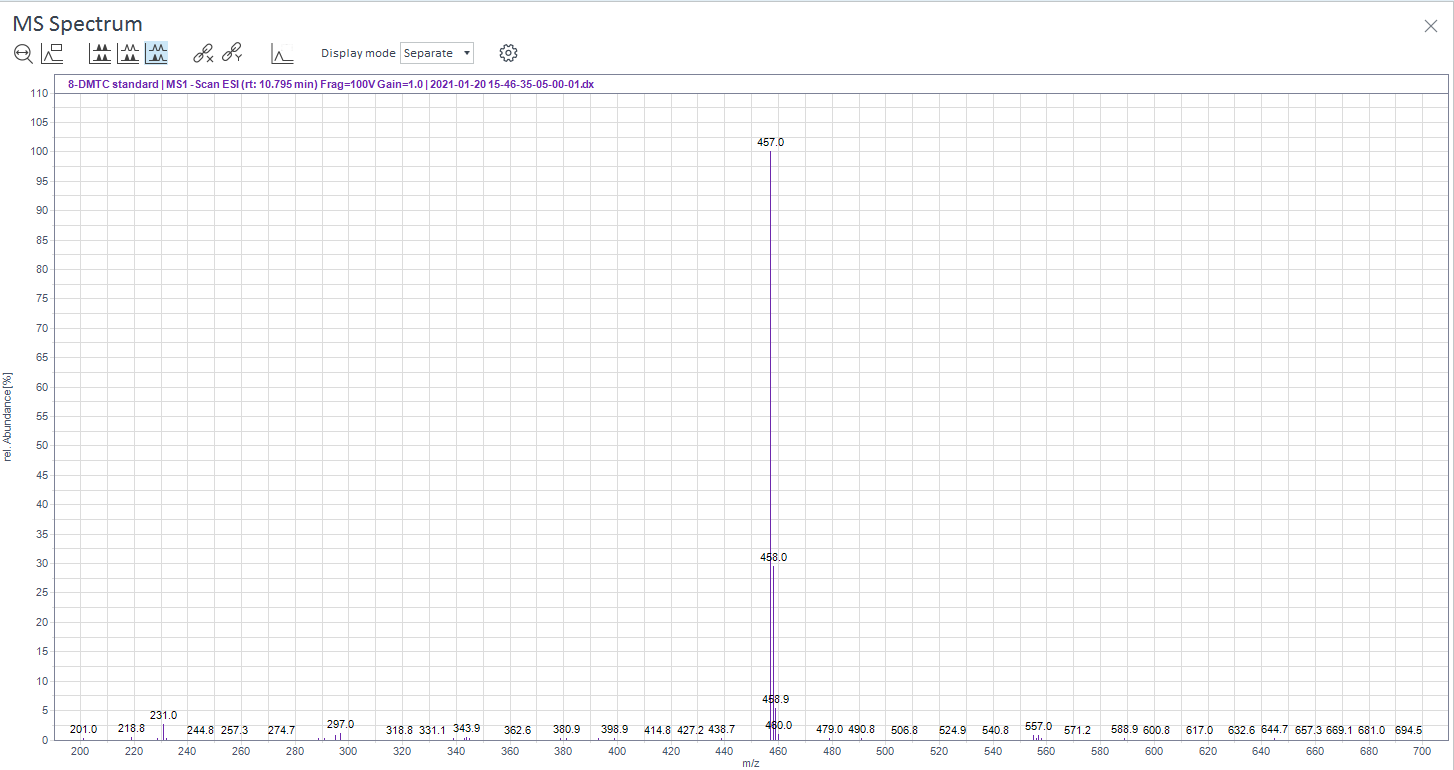

**Supplementary Figure 2** Mass spectrum for 8-demethyl-tetracenomycin C standard in ESI-MS negative ionization mode.

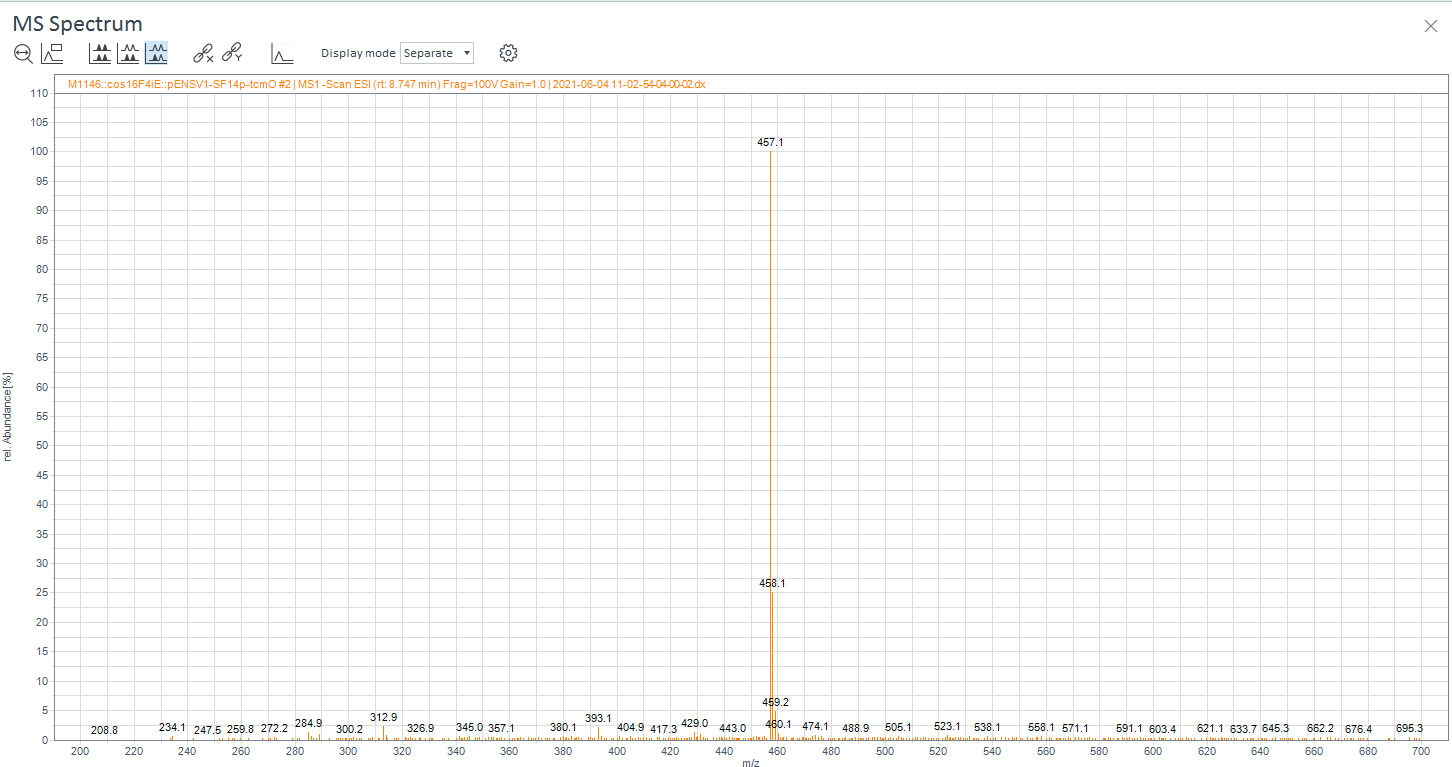

**Supplementary Figure 3** Mass spectrum for 8-demethyl-tetracenomycin C biosynthesized from a representative biological sample.

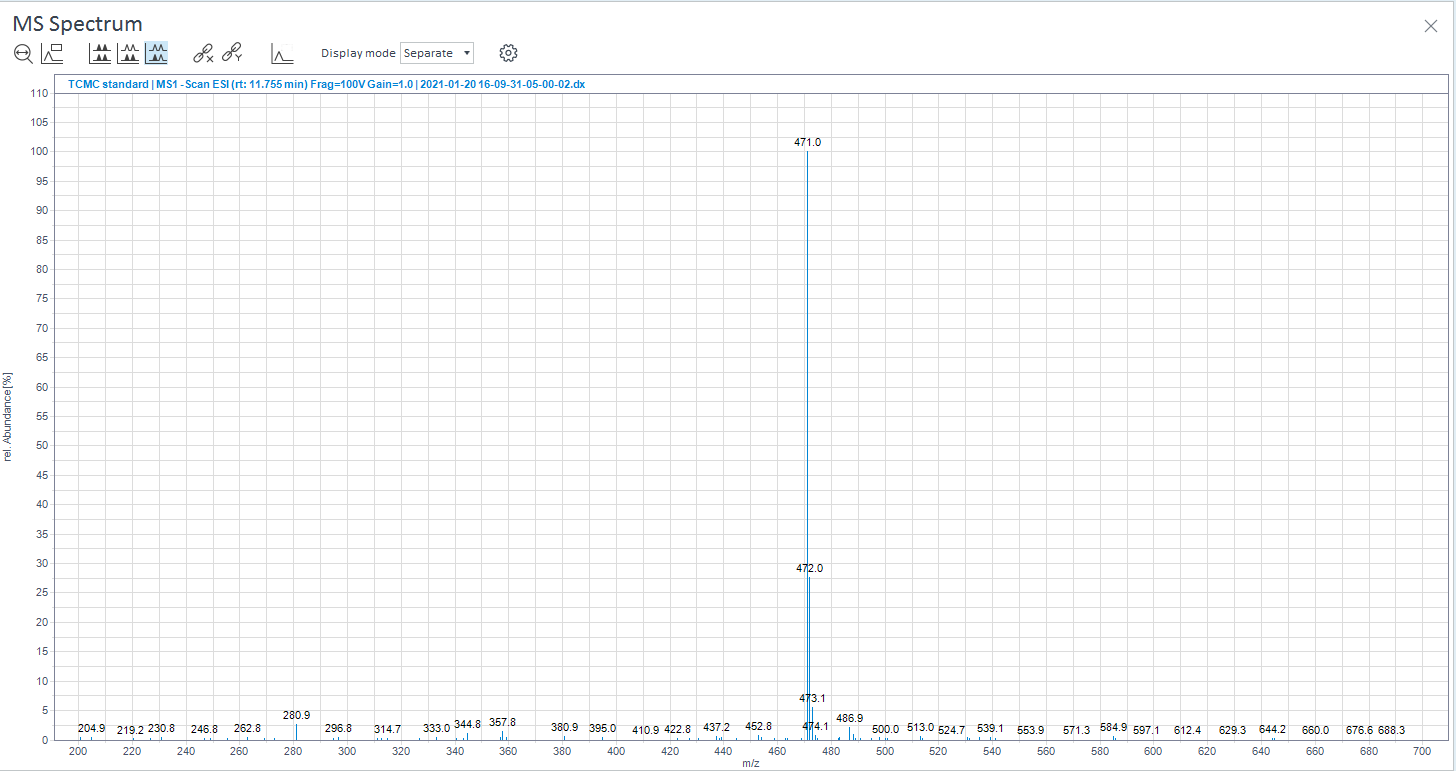

**Supplementary Figure 4** Mass spectrum for tetracenomycin C standard in ESI-MS negative ionization mode.

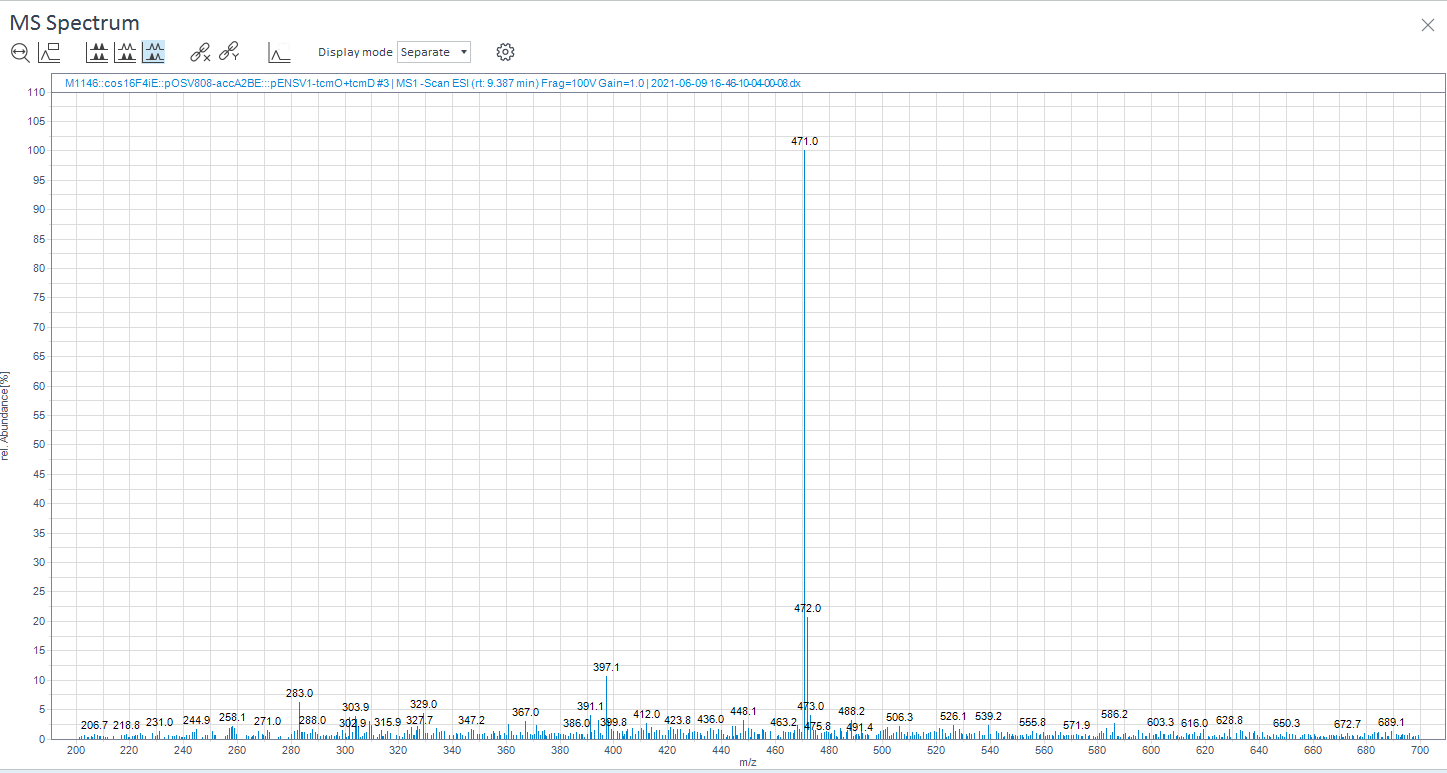

**Supplementary Figure 5** Mass spectrum for tetracenomycin C biosynthesized from a representative biological sample.

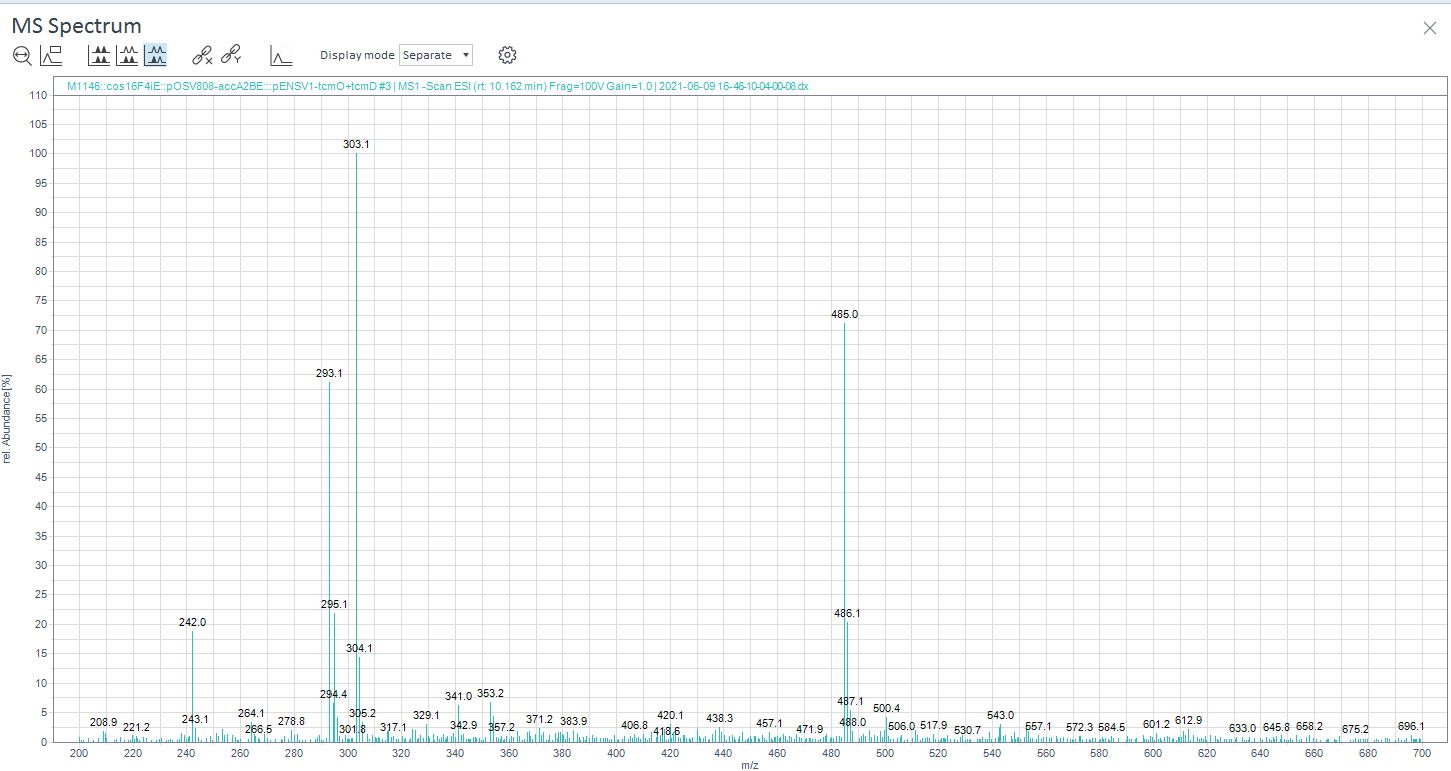

**Supplementary Figure 6** Mass spectrum for tetracenomycin X biosynthesized from a representative biological sample.

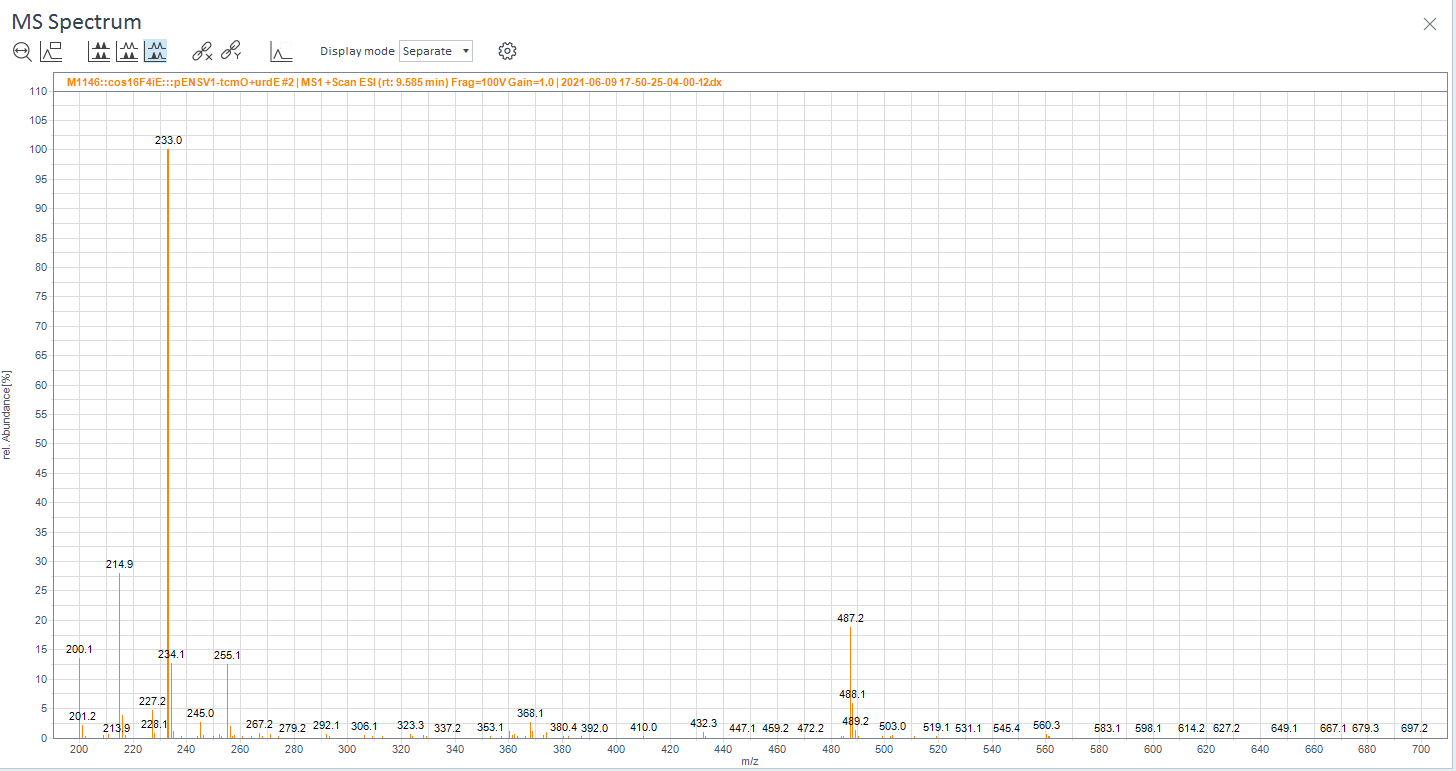

**Supplementary Figure 7** Mass spectrum for 6-hydroxy-tetracenomycin C standard in ESI-MS negative ionization mode.

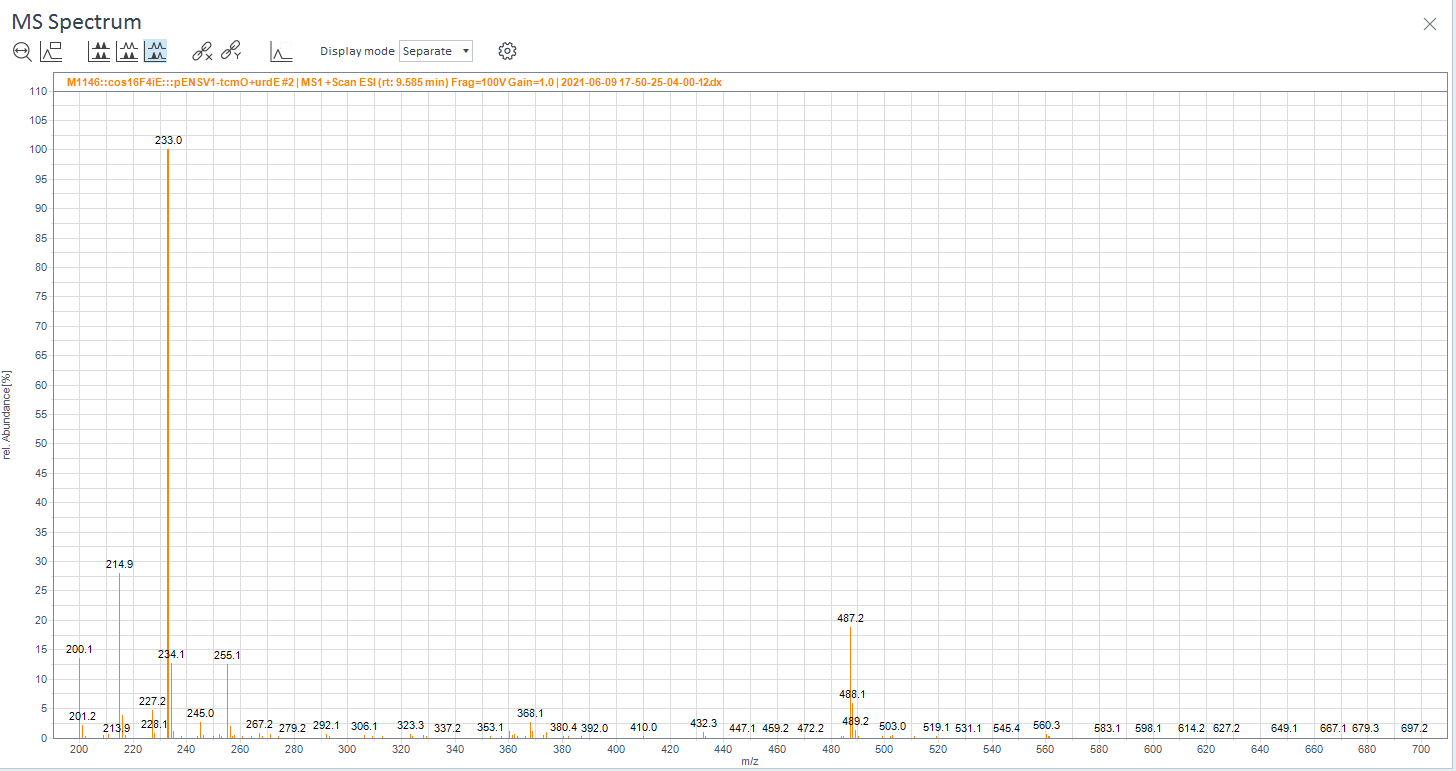

**Supplementary Figure 8** Mass spectrum for 6-hydroxy-tetracenomycin C biosynthesized from a representative biological sample.

References

Aubry, C., Pernodet, J.L., Lautru, S., 2019. Modular and integrative vectors for synthetic biology applications in Streptomyces spp. Applied and Environmental Microbiology 85. https://doi.org/10.1128/AEM.00485-19

Bentley, S.D., Chater, K.F., Cerdeño-Tárraga, A.-M.A.-M., Challis, G.L., Thomson, N.R., James, K.D., Harris, D.E., Quail, M.A., Kieser, H., Harper, D., Bateman, A., Brown, S., Chandra, G., Chen, C.W., Collins, M., Cronin, A., Fraser, A., Goble, A., Hidalgo, J., Hornsby, T., Howarth, S., Huang, C.-H., Kieser, T., Larke, L., Murphy, L., Oliver, K., O’Neil, S., Rabbinowitsch, E., Rajandream, M.-A., Rutherford, K., Rutter, S., Seeger, K., Saunders, D., Sharp, S., Squares, R., Squares, S., Taylor, K., Warren, T., Wietzorrek, A., Woodward, J., Barrell, B.G., Parkhill, J., Hopwood, D.A., 2002. Complete genome sequence of the model actinomycete Streptomyces coelicolor A3(2). Nature 417, 141–147. https://doi.org/10.1038/417141a

Bierman, M., Logan, R., O’Brien, K., Seno, E.T., Nagaraja Rao, R., Schoner, B.E., 1992. Plasmid cloning vectors for the conjugal transfer of DNA from Escherichia coli to Streptomyces spp. Gene 116, 43–49. https://doi.org/10.1016/0378-1119(92)90627-2

Flett, F., Mersinias, V., Smith, C.P., Flett ’, F., Mersinias, V., Smith, C.P., 1997. High efficiency intergeneric conjugal transfer of plasmid DNA from *Escherichia coli* to methyl DNA-restricting Streptomycetes. FEMS Microbiology Letters 155, 223–229. https://doi.org/10.1016/S0378-1097(97)00392-3

Yanisch-Perron, C., Vieira, J., Messing, J., n.d. Improved Ml3 phage cloning vectors and host strains: nucleotide sequences of the M13mp18 and pUC19 vectors (Recombinant DNA; molecular cloning; polycloning sites; progressive deletions).
